# Exploring the binding of resveratrol to an oncogene promoter DNA sequence d(CCAATTGG)_2_ through multispectroscopic, nuclear magnetic resonance and molecular dynamics studies

**DOI:** 10.1101/2020.09.10.291336

**Authors:** Shailendra Kumar, Peeyush Kumar, Maya S. Nair

## Abstract

We report the interaction of resveratrol with an octamer DNA sequence d(CCAATTGG)_2_, present in the promoter region of many oncogenes, using a combination of absorption, fluorescence, calorimetric and nuclear magnetic resonance techniques to probe the binding. Resveratrol binds to the duplex sequence with a binding constant 2.20×10^6^ M^−1^ in absorption studies. A ligand-duplex stoichiometry of 2.2:1 was obtained with binding constant varying from 10^9^ to 10^6^M^−1^ with the concentration of DNA varied in fluorescence titration measurements. Spectral changes indicated external binding of resveratrol to duplex DNA. Circular dichroism data displayed minimal variation suggesting external binding. Melting temperatures of DNA and its 1:1 complex showed a difference of approximately 2.25°C, which supports the external binding. Nuclear magnetic resonance data showed resveratrol binds to the minor groove region near the AT basepair from the nuclear Overhauser effect spectroscopic cross peaks. Distance restrained molecular dynamics was employed in explicit solvent condition to obtain the lowest energy structure. The complex was stable and retained the B-DNA conformation. Findings in this study identify resveratrol as a minor groove binder to the AT region of DNA and pave the way for exploring resveratrol and its analogues as promising anticancer/antibacterial drug.

## 1. Introduction

Deoxyribonucleic acid is one of the most significant molecular targets of therapeutic drugs and plays a significant role in various cellular processes such as replication, transcription and translation. During reproduction, DNA replication takes place and genetic information transferred to the next generation. This process is highly controlled at the molecular level. The new traits formed are governed by the gene sequences of DNA, which varies from healthy to fatal diseases [1,2]. Life-threatening conditions like cancer are a massive threat due to uncontrolled cell growth and cell division. Targeting DNA is a crucial step to stop the replication and transcription which control or inhibit the gene regulatory pathways in various diseases [3,4]. The DNA-drug interaction is of great interest in the various research fields of pharmacological and molecular medicine. Investigating the structural and functional characteristics of biomolecules can provide more insight into the interaction between the two [2]. It can help in understanding the mechanism of action and mode of association between the two. Often, this strategy is used to develop new DNA targeted therapeutic molecules [5]. Many natural and synthetic molecules with pharmacological properties show active binding to DNA [6,7].

Resveratrol (trans-3,4,5-trihydroxystilbene) is a natural stilbene derivative found in plants such as mulberries, peanuts, grapes and also found in plant parts like seeds, petioles, woody parts. It is a polyphenolic phytoalexin, present in many food products like red wine, grape juice etc. [8,9,10]. The basic structure of resveratrol comprises of two phenol rings connected through a double styrene bond. The double bond character is responsible for the isomeric state that exists in cis- and trans-form of resveratrol. Trans-form is the most stable conformation of resveratrol and is widely used for various therapeutics and also occurs in medical preparations [10,11]. It can regulate several pathways which are involved in cell proliferation, apoptosis and inflammation [12]. Various cell culture studies reveal that resveratrol can reduce cell cycle progression and can activate apoptotic cell death [9, 13–15]. Many researchers demonstrated that resveratrol induces the apoptotic activity of p53 [16,17].

Interaction of resveratrol with different DNA sequences for sequence-specific binding mode has been reported. Spectroscopic studies showed resveratrol binds to calf thymus DNA through intercalation. While docking studies showed intercalation as well as groove binding. It is also said that in intercalative mode, it is surrounded by ATTATT segment while in groove binding, the surrounding nucleotides are AATAAT [8]. Resonance light scattering techniques showed that resveratrol could be used for the detection of traces of DNA, based on its DNA binding property [9]. It has been demonstrated that resveratrol inhibited the growth of preneoplastic lesion in carcinogen-treated mouse mammary glands. Also, it hindered tumorigenesis in mouse skin cancer model [18]. Usha et al. showed that resveratrol could modulate DNA intercalation of dyes and protect the DNA from the damage induced due to the planar dyes, thereby reducing the DNA-directed toxicity [19]. Binding constants were calculated using absorption, and Fourier transform infrared (FTIR) spectroscopy for the drug-DNA and drug-RNA complexes. Partial stacking and H-bond formation with DNA/RNA is responsible for the antineoplastic activity of resveratrol [20].

Short sequences such as TATA, CCAAT, GC etc. are present in specific regions of promoters and enhancers which activates polymerase II enzyme transcribed mRNA genes. Few such stretches like TATA, GC and CCAAT boxes occur at extremely high frequency [21]. The CCAAT box was the first element identified [22]. Later studies established that such pentanucleotide sequences are present in a wide variety of vertebrate, yeast and plant promoters and are essential for transcription. All five nucleotides are almost invariably conserved. CCAAT sequences can be found both in the direct and in the inverted orientation, and it is present in both TATA containing and in TATA-less promoters. Statistical analysis of 500 unrelated proteins revealed that CCAAT pentanucleotide is present in 30% of them. Many sequence-specific regulators recognize these short nucleotides. NF-Y is a nuclear protein which recognizes the pentanucleotide CCAAT. The binding site of CCAAT-enhancer binding protein (c/EBP) contains the CCAAT nucleotides in the intervening sequences. CCAAT transcription factor (CTF/NF-1) binds to viral and cellular promoters, recognize sequences containing CCAA nucleotides [15,23]. Borghini et al. [24] pointed out the lack of conventional TATA boxes and the presence of two identical CCAAT boxes as the crucial elements involved in the transcription regulation of the human *TLX3* gene which is overexpressed in T-cell Acute Lymphocytic leukaemia.

The palindromic CCAATTGG sequence is present in promoters of human, fly and rat genomes which regulate the activity of several growth factors, oncogenes and tumor suppressor genes. Fibulin-5 gene (FBLN5) promoter contains palindromic sequence CCAATTGG at upstream of the first translation initiation codon, which functions as a potential CCAAT/enhancer binding element [25]. The CCAATTGG motif occurs in BEN (**B**ANP, **E**5R, and **N**AC1) domain promoter of drosophila insensitive (Insv) gene which is responsible for its neural development and is widely distributed in metazoans and viruses [26]. The mammalian BEN5/6, RBB and BTB domain protein also recognize a CCAATTGG palindromic motif with homologous Insvfactor[26–29]. The dyad-symmetric CCAAT core sequence is found in the CCAATTGG motif of rat argininosuccinatelyase (AL) promoter at the position of −80 base pair which involves in arginine biosynthesis and expressed in a high level in the liver [30]. Nuclear Factor-Y (NF-Y) similar to transcription factor contains two subunits that show good affinity towards CCAAT central motif in several promoters. Point mutations or spaces in the half-site of this motif inhibit the binding of NF-Y to CCAATTGG palindromic sites. This finding establishes that NF-Y recognizes the correct reading frame dyad symmetric CCAAT core motif [31]. Literature reveals that the CCAAT sequence is vital in many cellular processes that involve cell proliferation and cell integrity [32–35].

Different groups have studied the interaction between DNA sequences comprising CCAAT segment with small molecules. In a sequence-specific interaction study, distamycin was found to bind at the central portion in the minor groove of duplex DNA d(GGCCAATTGG)_2_ [36]. Crystallographic analysis of distamycin-d(CGCAAATTTGCG)_2_complex showed the preference of distamycin to the AT rich region, in the minor groove of DNA [37]. Another molecule netropsin displayed a strong affinity to the central CAATTG segment of the sequence d(GGCCAATTGG)_2_ in the crystal structure [38]. NMR spectroscopic study on the binding of Hoechst 33258 revealed that two drug molecules bind independently at the 5’-TTTT and 5’-AAAA region of d(CTTTTGCAAAAG)2 duplex DNA sequence with interactions limited to four A.T base pairs [39]. Crystallographic studies on the interaction of Hoechst 33258 with sequences d(CGCGAATTCGCG)2 and d(CGCGATATCGCG)2 demonstrate that Hoechst molecule binds with central AT rich regions of the duplex DNA through hydrogen bonding, electrostatic interactions and van der Waal interactions [40–42].

Neidle’s group studied the sequence-specific binding of Hoechst with dodecamer duplex DNA (CGCAAATTTGCG)2 sequence in which Hoechst molecule recognizes the central ATTTG base pair patch via minor groove binding in specific orientations, with high affinity [43]. Crystal structure of berenil with dodecamer DNA sequence d(CGCAAATTTGCG)2 showed that the drug molecule binds to the 5’-AATT segment with hydrogen bonds formed between amidinium group of berenil and thymine bases of DNA [44]. Berenil also forms a complex with another dodecamer DNA sequence d(CGCGAATTCGCG)2. It binds to the region 5’-AAT forming hydrogen bonds to adenine atoms [45]. These studies highlight the sequence-specific binding of small molecules to DNA.

In this paper, we report the interaction of DNA sequence d-(CCAATTGG)_2_ with resveratrol to throw light on the binding mode and conformational changes to DNA upon binding. The DNA sequence d-(CCAATTGG)_2_ was selected because of two reasons: firstly, due to its biological significance as a promoter site in various oncogenes [23,46] and secondly due to its AT rich regions, which are strongly over-represented in origins of replication in many organisms [47]. We tried to investigate the binding site and structural changes that took place in the DNA sequence d-(CCAATTGG)_2_ upon interaction with resveratrol through absorption, fluorescence, circular dichroism (CD), differential scanning calorimetry (DSC) and nuclear magnetic resonance (NMR)studies. NMR derived distance parameters are employed to perform restrained molecular dynamics (MD) along with simulated annealing to obtain the final structural parameters of the DNA-resveratrol complex. Figure 1 represents the structure of resveratrol (A) and DNA sequence (B).

**Fig. 1.**
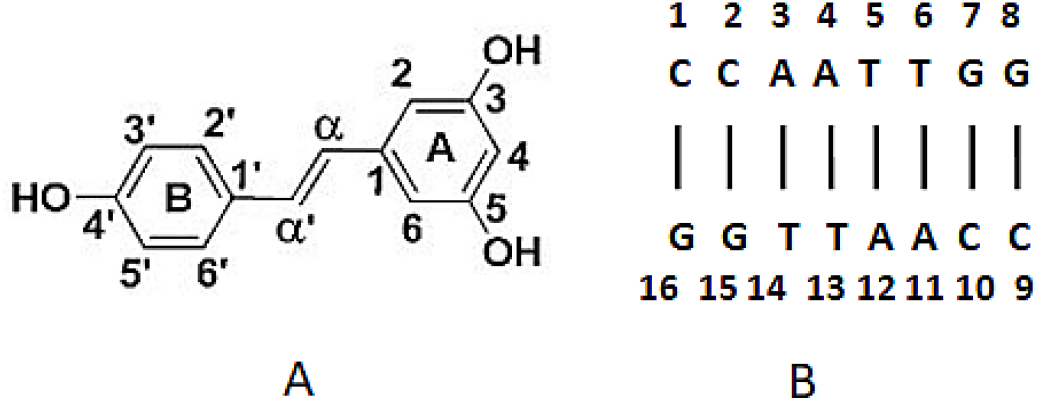
Schematic representation of (A) resveratrol and (B) d(CCAATTGG)_2_DNA sequence.

## 2. Materials and Methods

The single-stranded oligonucleotide sequence CCAATTGG and resveratrol were purchased from Sigma Aldrich Co., USA. DNA stock solution was prepared in 20 mM sodium phosphate buffer containing 10 mM sodium chloride and 0.2mM EDTA at pH 7.2. The DNA solution was heated and kept at 95°C for 5 minutes and annealed overnight to facilitate the formation of doublestranded DNA structure. Resveratrol was dissolved in 1:3 methanol-water. Concentrations of DNA and resveratrol were determined spectrophotometricallyusing molar extinction coefficients 76300 M^−1^ cm^−1^at 260 nm and 32,000 M^−1^ cm^−1^ at 318 nm respectively. All chemicals used were of analytical grade. Millipore water was used throughout the experiments.

### 2.1. UV–Visible absorption measurements

The absorption spectra of free DNA and complexes with resveratrol were carried out using CARY 60 spectrophotometer (Agilent Technologies). Absorption was recorded from 200 to 700 nm using a 1cm pathlength quartz cuvette. Spectra were collected with different concentrations of DNA (N) added to a fixed amount of resveratrol (D). Appropriate blank solutions were used for baseline subtraction. Binding coefficient K_b_ was calculated by using a double reciprocal plot as given below[48,49].

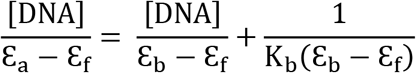

where Ԑ_f_, Ԑ_b_ and Ԑ_a_ are molecular extinction coefficients of the free, bound and apparent (observed) state respectively and [DNA] is the concentration of DNA.

### 2.2. Fluorescence measurements

Fluorescence experiments were performed using FluoroMax Plusspectrofluorometer(Horiba Scientific,USA). Fluorescence titrations were carried out in quartz cuvette with a fixed concentration of resveratrol at 10 μm. DNA solutionsweregradually addedin various D/Nratiosat room temperature. The emission spectra were recorded from 330 nm to 600 nm, with excitation wavelength at 318 nm. The excitation and emission slits were kept at 5 nm. Binding constant (K_b_) and the number of binding sites (n) were calculated from the double logarithmic plot using the equation[50].

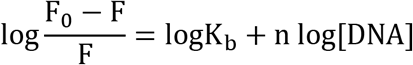

WhereF_o_ and F are the fluorescence intensities of the fluorophore in the absence and presence of different concentrations of DNA, respectively.

### 2.3. Circular Dichroism measurements

Circular dichroism experiments were carried out by using Chirascan CD spectrometer (Applied Photophysics, Leatherhead, UK) attached with a temperature controlled cell holder. Quartz cuvette of 0.1 cm path length was used for the measurements. CD spectra were recorded in the wavelength a range of 190 to 700 nm. The concentration of the DNA sequence was kept constant at 100μM during the titrations and resveratrol solutions were added serially in various D/N ratios at 25°C in phosphate buffer at pH 7.2. Spectra were averaged over three scans, and the baseline was subtracted.All CD spectra were processed using Chirascan software.

### 2.4. Differential Scanning Calorimetry

The Differential Scanning Calorimetry (DSC) experiments were carried out using a MicroCal VP-DSC instrument (MicroCal, Northampton, MA, USA). In this method, excess heat capacity as a function of temperature was measured to observe the thermal transitions from folded to the unfolded state. The DNA sample was prepared in 20 mM of sodium phosphate buffer. All samples were degassed before the experiment. Each sample was scanned from 25°C to 120°C with a scan rate of 60°C/h at constant pressure 33 psi. The reference and sample cells were loaded with the buffer solution and repeatedly scanned until proper baseline was obtained. 100 μM free DNA solution was loaded in the sample cell and scanned to obtain the thermal profile of free DNA. The complexes of DNA (N) with resveratrol (D) at D/N = 1 and 2 were scanned to obtain the thermal profile of bound forms. All scans were subtracted from the baseline, and thermograms were analyzed using inbuilt Origin software. Thenon-two-state model curve fitting was applied to fit the raw data of each thermogram.

All spectroscopic data were analyzed and plotted using Origin software.

### 2.5. Nuclear Magnetic Resonance experiments

NMR samples of DNA were prepared by dissolving dry oligonucleotide to give 3.4 mM solution in 90% H_2_O and 10% D_2_O containing 10 mM sodium phosphate buffer, pH 7, 0.25 mM EDTA and 10 mM sodium chloride. Samples of resveratrol (28 mM) were prepared by dissolving the compound in d-ethanol. DNA-drug complexes at different Drug (D) to Nucleic acid (N) duplex ratio (D/N) were prepared by titrating different concentrations of DNA and drug. Trimethylsilyl propionic acid (TSP) was used as an internal reference.

All NMR data were collected on a Bruker Avance AMX-500 MHz FT-NMR spectrometer located at Nuclear Magnetic Resonance (NMR) Facility, Indian Institute of Technology Roorkee. The data obtained were processed with TOPSPIN version 1.3 software (Bruker). One dimensional (1D) NMR were collected at 278 K, 283 Kand 298 K with 64 K data points, 64-128 number of scans, spectral width of 12/20 ppm. The removal of HOD signal was accomplished using watergate suppression. The two dimensional (2D) phasesensitive nuclear Ovehauser Effect spectroscopy (NOESY) experiments were carried out at 283 K and 298 K with mixing times τ_m_ = 200 and 300 ms.Typical parameters for 2D experiments were: 2048 data points along *t*_2_ dimension; 512 free induction decays in *t*_1_ dimension; pulse width ≈ 9.5–12 μs; spectral width ≈ 5000 Hz; no. of scans = 64–128; digital resolution 2.30–4.60 Hz/point and relaxation delay ≈ 2.0 s. Hetero Nuclear Multiple Bonded Correlation spectroscopy (HMBC),TOtal Correlation SpectroscopY (TOCSY) were carried out at 283 K with 400/512 by 2048 data points. One dimensional^31^P and Two dimensional phosphorous exchange (^31^P-^31^P) spectra were collected using mixing times of 150 ms and 200 ms for different D/N ratios at 283 K and 298 K.

Distance restraints were obtained from the volume of experimental NOE cross peaks, calibrated with cytosine H5-H6, using a distance of 2.45 Å in Sparky software[51].A range of ±0.5 Å was provided to avoid any errors in integration.

Distance restrained Molecular Dynamics (rMD) was performed using AMBER 18 simulation package [52]and the protocols used were described in the relevant section.

All helical parameters, backbone torsional angles, and sugar conformations of the resulting rMD structures were analyzed using Curves+software[53].

## 3. Results and Discussion

### 3.1. Absorption studies

The absorption spectrum of resveratrol showed a broad absorption spectrum with two kinks around 304 nm and 318 nm. The titration was carried out with a fixed concentration of resveratrol. The gradual addition of oligonucleotide d(CCAATTGG)_2_ to resveratrol (N/D) in a set of 25 experiments resulted in a sequential hyperchromic effect without any noticeable shift in the maximum absorbance wavelength as shown in Fig. 2A. Data were analyzed with absorption at 318 nm. A plot of absorbance at 318 nm against DNA concentration shows a linearly increasing curve until saturation around N/D = 2 (Fig. 2B). The binding constant was calculated using the double reciprocal plot (Fig. 3) obtained as K_b_ = 2.20 ×10^6^ M^−1^. Earlier our studies showed that resveratrol binds to d(CGAATTCG)2 and d(CGTTAACG)2 sequences with binding constants of the order of 10^5^ [54]. The results of the present study, therefore indicate that resveratrol binds more strongly to the sequence CCAATTGG as compared to the above two sequences, pointing to the sequence specificity of the compound. Absence of any isosbestic point suggests that either no equilibrium state or multiple equilibrium states exist in the solution. Hyperchromism with no shift in absorption maxima usually results from an external binding of the molecule to the DNA sequence. Often hyperchromism with a bathochromic shift indicates an intercalative mode of binding. The observed hyperchromism with no change in absorbance maxima, therefore, indicate the external binding of resveratrol.

**Fig. 2.**
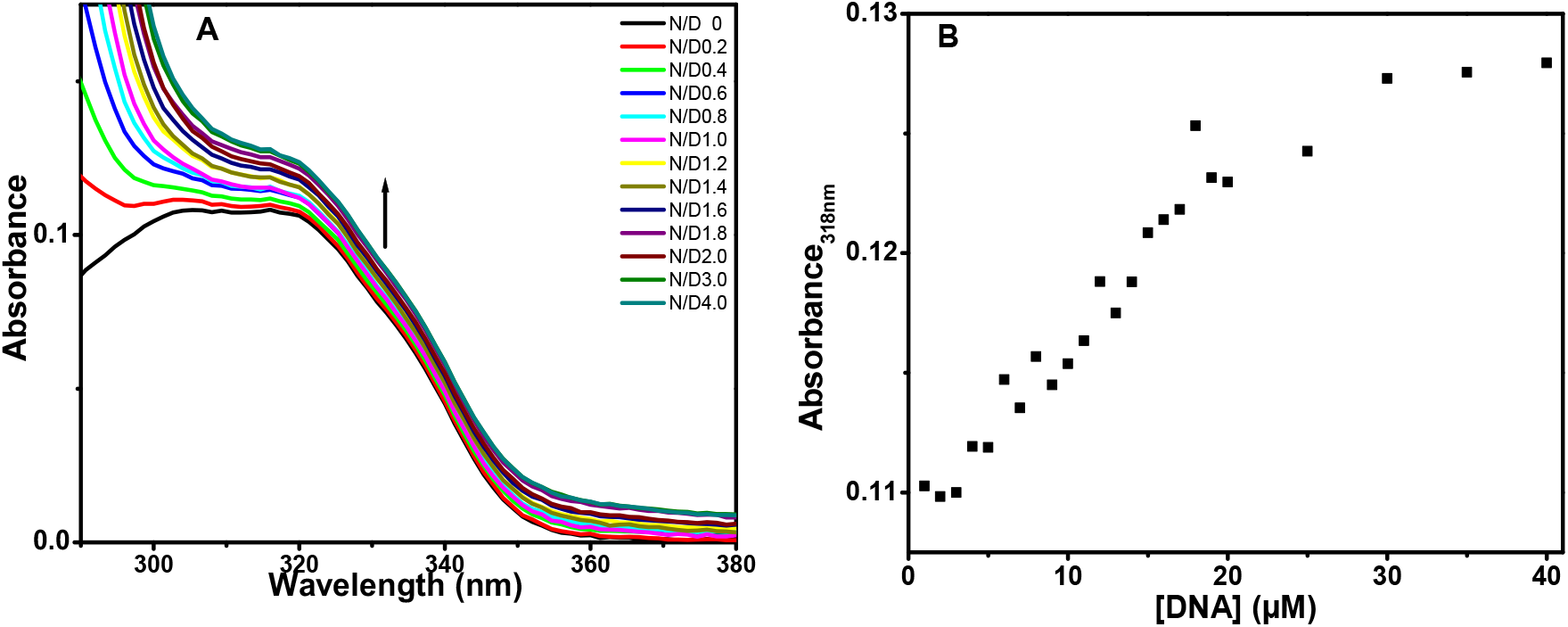
Absorption spectra of resveratrol in the presence of different concentrations of DNA (A) N/D =0 to 4, (B) Variation in absorbance of resveratrol with DNA concentration

**Fig.3.**
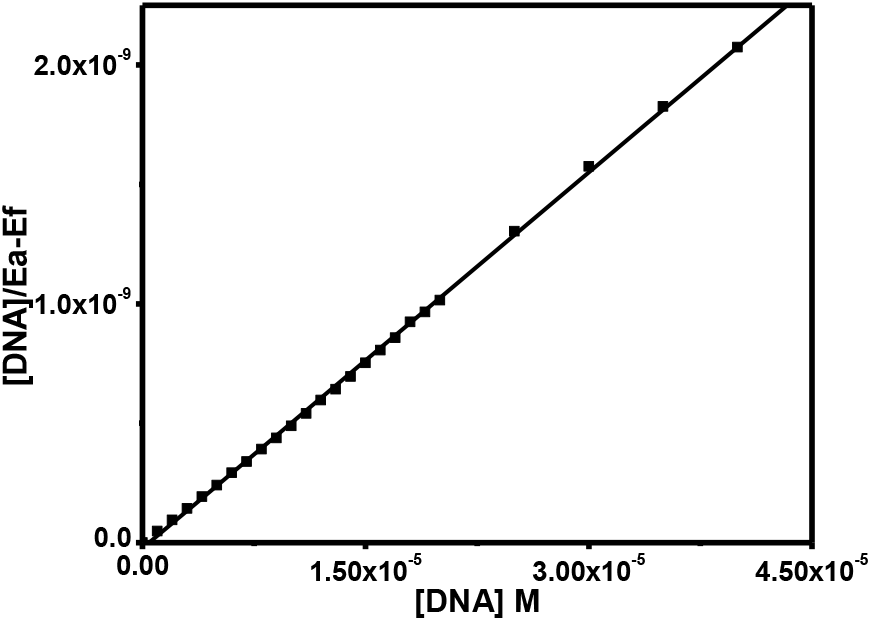
Calculation of binding constant from [DNA]/(Ea-Ef) vs [DNA] curve.

### 3.2. Fluorescence studies

Resveratrol gives intense emission around 388 nm upon excited at 318 nm. The broadband is peaked at 381nm with two shoulders at 353 nm and 410 nm. These peaks are attributed to the emission from a noncharged form of the resveratrol as described elsewhere[55]. Fig. 4 illustrates the fluorescence emission spectra of resveratrol alone and in the presence of DNA at different nucleic acid to ligand ratios (N/D). Incremental addition of DNA to a fixed concentration of resveratrol resulted in a blue shift, from 388 nm to 381nm. At the same time, the 353 nm band red shifted to 364 nm along with an increase in emission intensity. The spectral changes indicate the interaction between DNA and resveratrol. Resveratrol was added to DNA till saturation is reached, around N/D = 3.0. The differences suggest that the hydroxyl groups of the ligand are in different environments [55]. In general, when the ligand is in a hydrophobic environment, its fluorescence intensity increases [56]. The fluorescence intensity almost doubled at N/D = 1and increased by four times when DNA concentration is 20 μM (N/D = 2) (Fig. 5). Such kind of increase in fluorescence when a ligand binds to AT rich DNA has been reported. This can be due to the lowering of the conformational flexibility of the compound within the complex [57,58]. This property of the molecule can be exploited to develop DNA sequence selective probe, and our group is working to this goal.

**Fig. 4.**
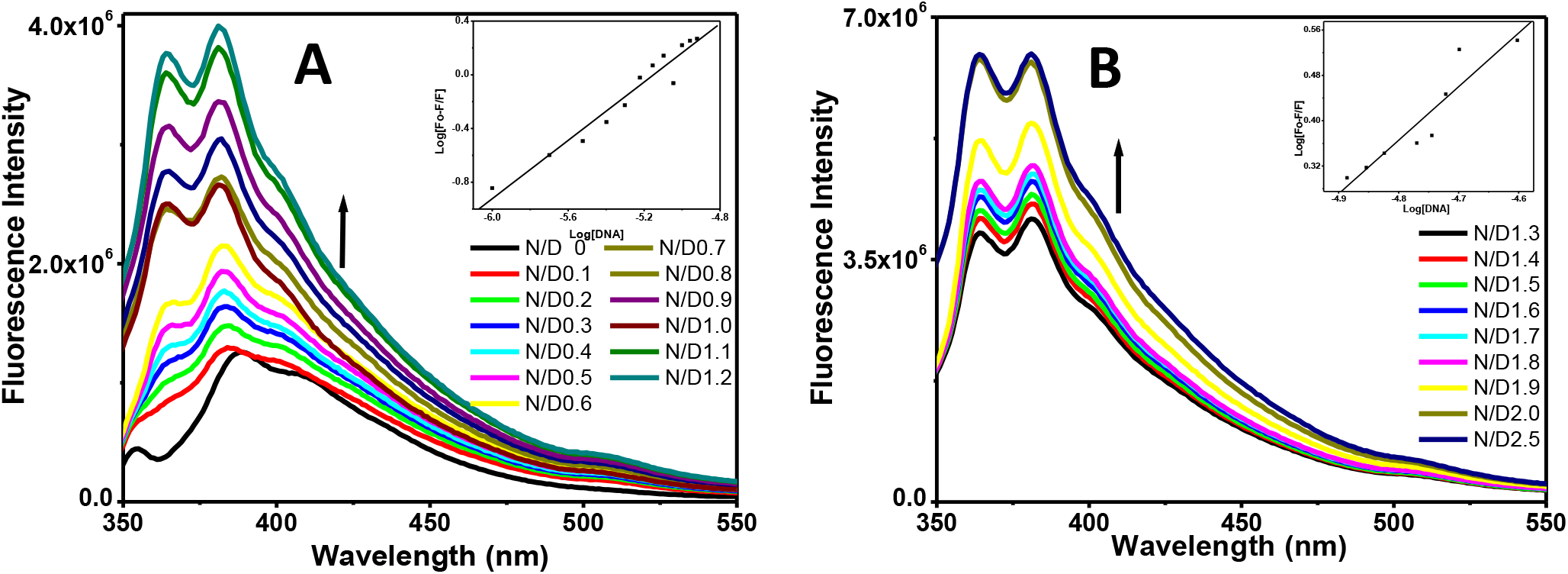
Fluorescence spectra of resveratrol with different concentrations of DNA, (A) N/D ratio = 0 to 1.4 and B) N/D ratio =1.4 to 2.3. Inset shows the plot for the calculation of binding constant.

**Fig. 5.**
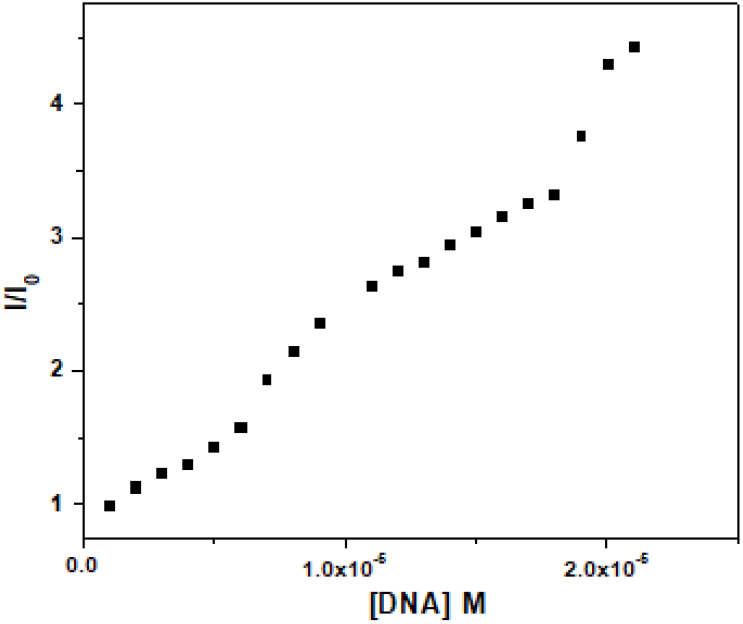
Relative fluorescence intensities of resveratrol as a function of DNA concentration.

The Binding constant (K_b_) and the number of binding sites (n) were calculated using double logarithmic plot (inset of Fig. 4A) and obtained as 7.94×10^9^ M^−1^ and 1.8 respectively for N/D = 0 to 1.4. Whereas for N/D= 1.4 to 2.3, K_b_ was obtained as 8.0×10^4^ M^−1^) and n = 0.76 (Inset of Fig. 4B) respectively. N=2.3 indicates that there are at least two strong binding sites in the DNA sequence. The binding stoichiometry of the complex was measured using the Job plot (Continuous variation method). The total concentration of ligand and DNA was kept constant at 20 μM. The emission intensity of resveratrol at 388 nm was plotted against the mole fraction of resveratrol as shown in Fig. 6. The cross over point was obtained at mole fraction equal to 0.69 which indicates a stoichiometry (χ / 1-χ, where χ is the mole fraction of ligand) of 2.2: 1 indicating that two molecules of resveratrol bind with one molecule of DNA.

**Fig. 6.**
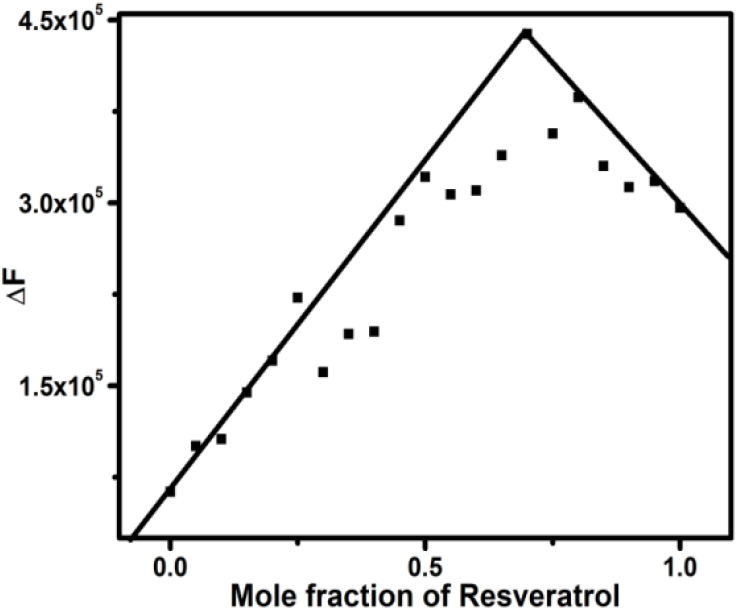
Job plot for the binding of resveratrol to DNA. The difference in fluorescence intensity at 388 nm as a function of mole fraction of resveratrol was plotted.

### 3.3. Circular dichroism spectroscopy (CD) studies

Circular dichroism (CD) spectroscopy is an important technique which displays structural changes in DNA sequences. The CD spectrum of canonical right handed B-DNA exhibits a positive band around 269 nm, owing to base stacking and a negative band around 247 nm, owing to helicity [59]. Often changes in the CD signals of DNA on interaction with ligand is attributed to the corresponding changes in DNA structure. Resveratrol did not show any CD band. The changes in CD spectrum of DNA were monitored upon ligand binding, which is due to corresponding changes in DNA conformations.

Fig. 7 describes the variation in CD bands when different concentrations of resveratrol were added to DNA (D/N ratio). The intensities of the positive band around 269 nm and negative band around 247 nm decreased till D/N = 5.0 without any noticeable shift. The 269 nm band decreased by 40%, while the 247 nm band decreased by 22%, indicating that the interaction between resveratrol with DNA caused a disturbance in the stacking of the bases as well as helicity. Absence of any significant spectral shift and induced CD band rules out the possibility of intercalation. The spectral changes observed suggest that resveratrol is binding externally and induce slight variations in the conformation of DNA.

**Fig. 7.**
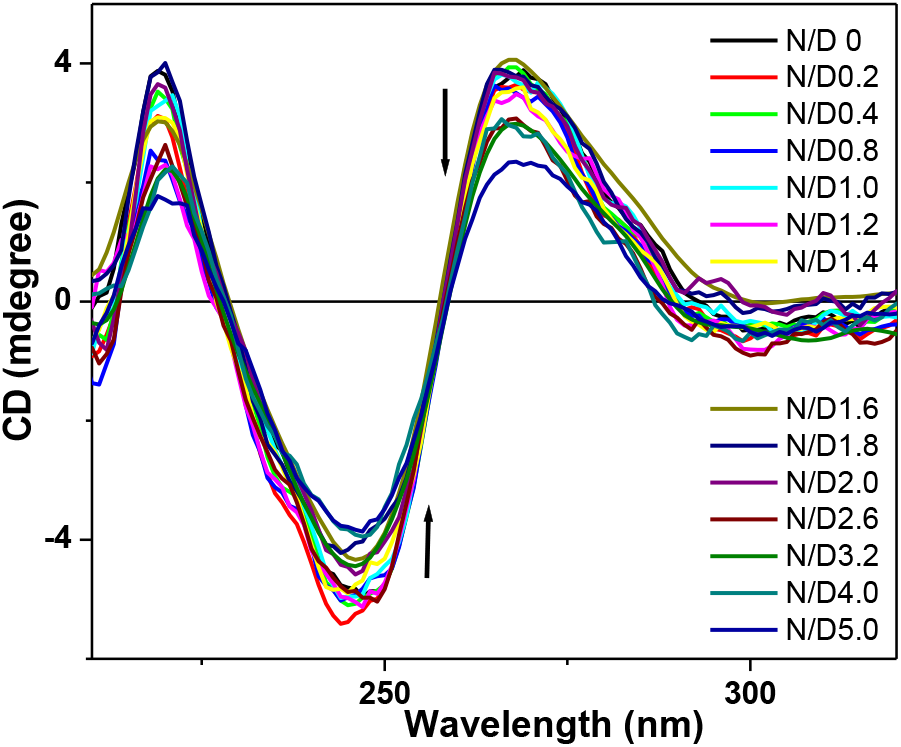
CD spectra of DNA alone (100 μM) and with different concentration of resveratrol.

### 3.4. Differential Scanning Calorimetry

Melting temperature (Tm) due to transition from complexed to uncomplexed (ordered to disordered) state can be estimated by evaluating the excess heat capacity as a function of temperature using differential scanning calorimeter (DSC). DSC thermograms were obtained for alone DNA and its complex with resveratrol at N/D =1 and 2 as presented in Fig. 8. In the case of alone DNA, the thermogram gave a good fit with two peaks, one major peak centred at T_m1_= 39.25°C and another peak around T_m2_ = 46.27°C. More than one T_m1_ shows that the melting pathway consists of intermediate species[60]. At N/D = 1.0, T_m1_ is shifted to 42.20°C and T_m2_ to 5I.23°C. ΔT_m1_ = 2.95°C observed from changing N/D = 0 to1 indicates a minimal change, pointing to the external binding of the compound to DNA. This result correlates with the observations of absorption and fluorescence findings. At N/D = 2.0, the DSC thermogram fits with two peaks with T_m1_ = 42.O7°C and T_m2_ = 52.92°C as indicated in Table 1. Positive (n) of van’t Hoff enthalpy to calorimetric enthalpy gives an estimate of the effective number of base pairs in a cooperative melting system[62,63]. Apparent values of n show that at least 6 base pairs are there in the cooperative melting system.

**Fig. 8.**
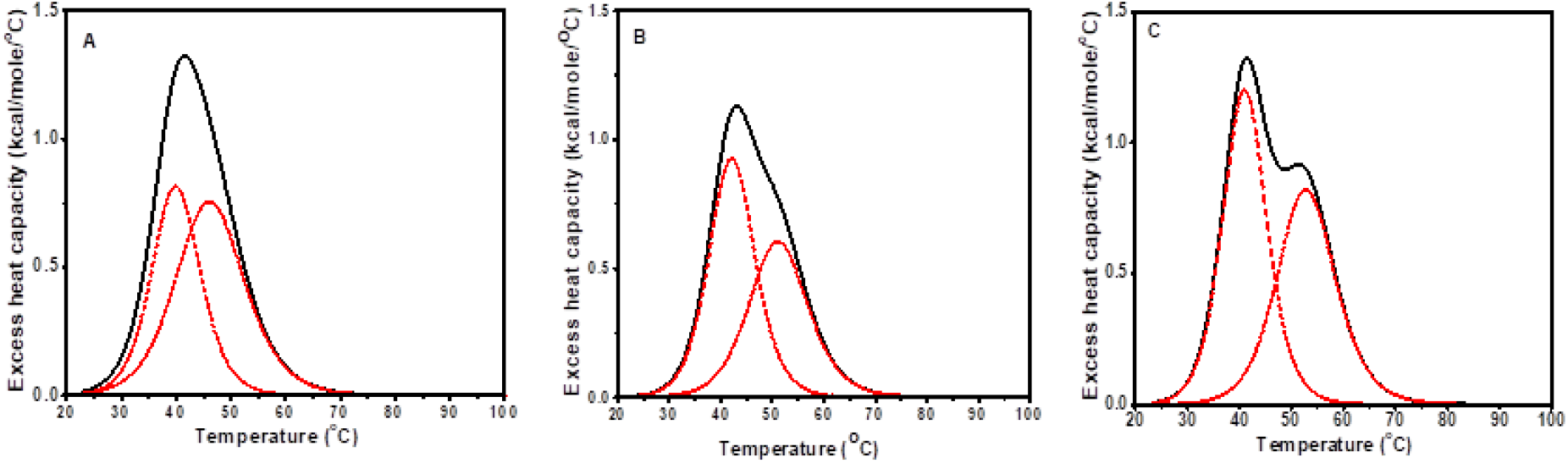
Differential Scanning Calorimetry curves and fitted data for A) alone DNA, B) DNA: resveratrol 1:1 complex: and C) 1:2 complex.

**Table 1.**
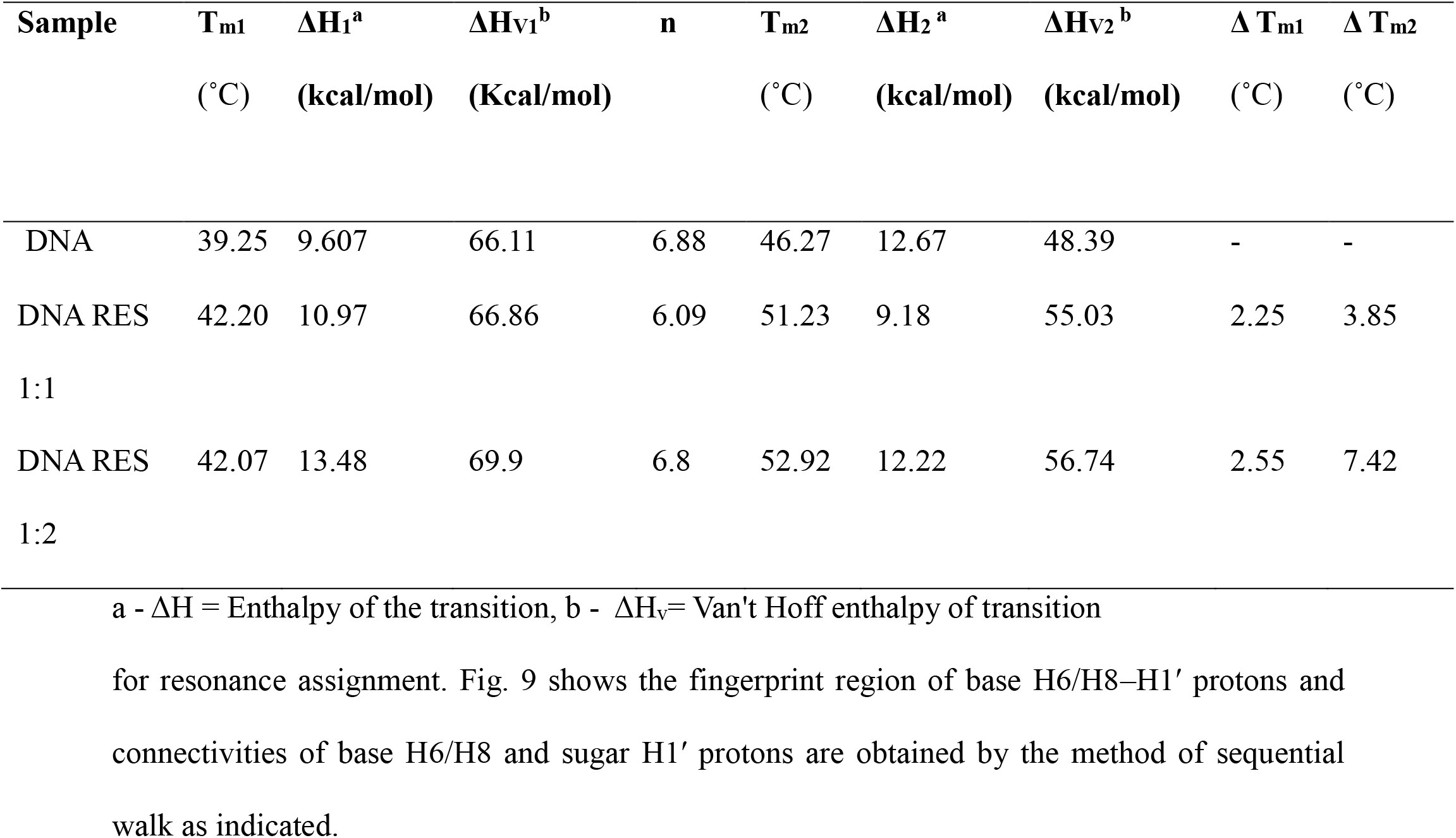
Thermodynamic parameters obtained from DSC

| Sample | T<sub>m1</sub> | ΔH<sub>1</sub><sup>a</sup> | ΔH<sub>v1</sub><sup>b</sup> | n | T<sub>m2</sub> | ΔH<sub>2</sub><sup>a</sup> | ΔH<sub>v2</sub><sup>b</sup> | Δ T<sub>m1</sub> | Δ T<sub>m2</sub> |
| --- | --- | --- | --- | --- | --- | --- | --- | --- | --- |
|  | (°C) | (kcal/mol) | (Kcal/mol) |  | (°C) | (kcal/mol) | (kcal/mol) | (°C) | (°C) |
| DNA | 39.25 | 9.607 | 66.11 | 6.88 | 46.27 | 12.67 | 48.39 | - | - |
| DNA RES<br>1:1 | 42.20 | 10.97 | 66.86 | 6.09 | 51.23 | 9.18 | 55.03 | 2.25 | 3.85 |
| DNA RES<br>1:2 | 42.07 | 13.48 | 69.9 | 6.8 | 52.92 | 12.22 | 56.74 | 2.55 | 7.42 |

a - ΔH = Enthalpy of the transition, b - ΔH<sub>v</sub>= Van't Hoff enthalpy of transition

### 3.5 Proton Resonance Assignments

#### 3.5.1. d - (CCAATTGG)_2_

One dimensional proton spectra were recorded for the sample containing DNA at 278K, 283K and 298K in 90% H2O and 10% D2O. The NMR resonances of the two strands of the octamer coincide due to the 2-fold symmetry of the duplex, and only one set of resonances was observed. The assignment of nucleotide protons (Supplementary material Table 1A) was carried out following the strategies available in the literature for sequential assignment in right handed B-DNA [64–66]. NOESY spectrum of DNA recorded at 298K with a mixing time 200ms was used

#### 3.5.2. Resveratrol

The protons of resveratrol have been assigned with the help of 1D proton NMR, Rotating Frame Overhauser Effect Spectroscopy (ROESY) and Total Correlation Spectroscopy (TOCSY) experiments (not shown). The chemical shifts of resveratrol were obtained, as shown in supplementary material Table 2. The one dimensional proton spectra of resveratrol in deuterated ethanol and deuterium oxide (D_2_O) at 298K are shown in Fig. 10. The results were consistent with the literature on the NMRstudies of resveratrol[67,68]

**Fig. 9.**
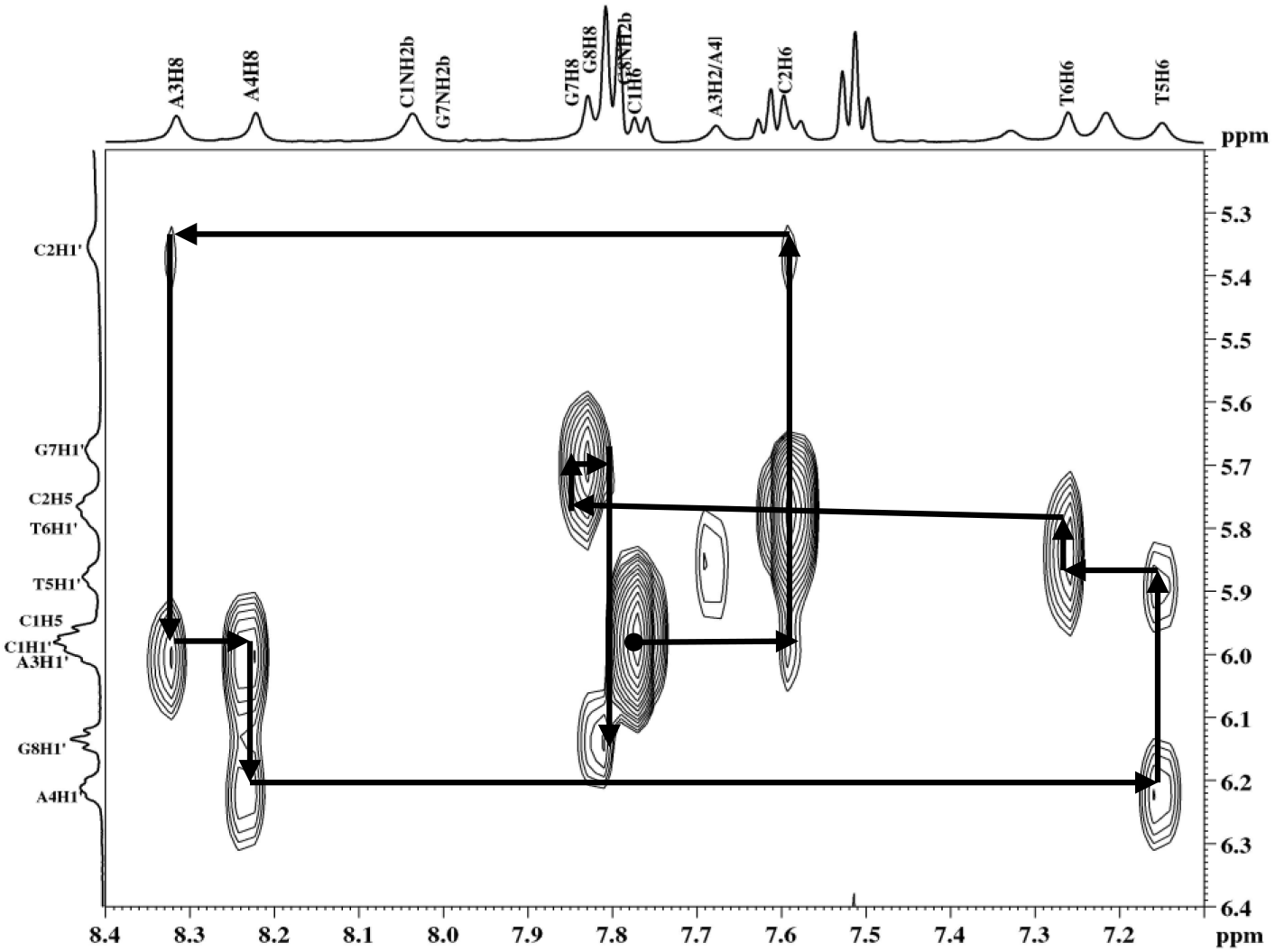
NOESY spectrum of d(CCAATTGG)_2_showing base H6/H8 – sugar H1′ connectivities.

**Fig. 10.**
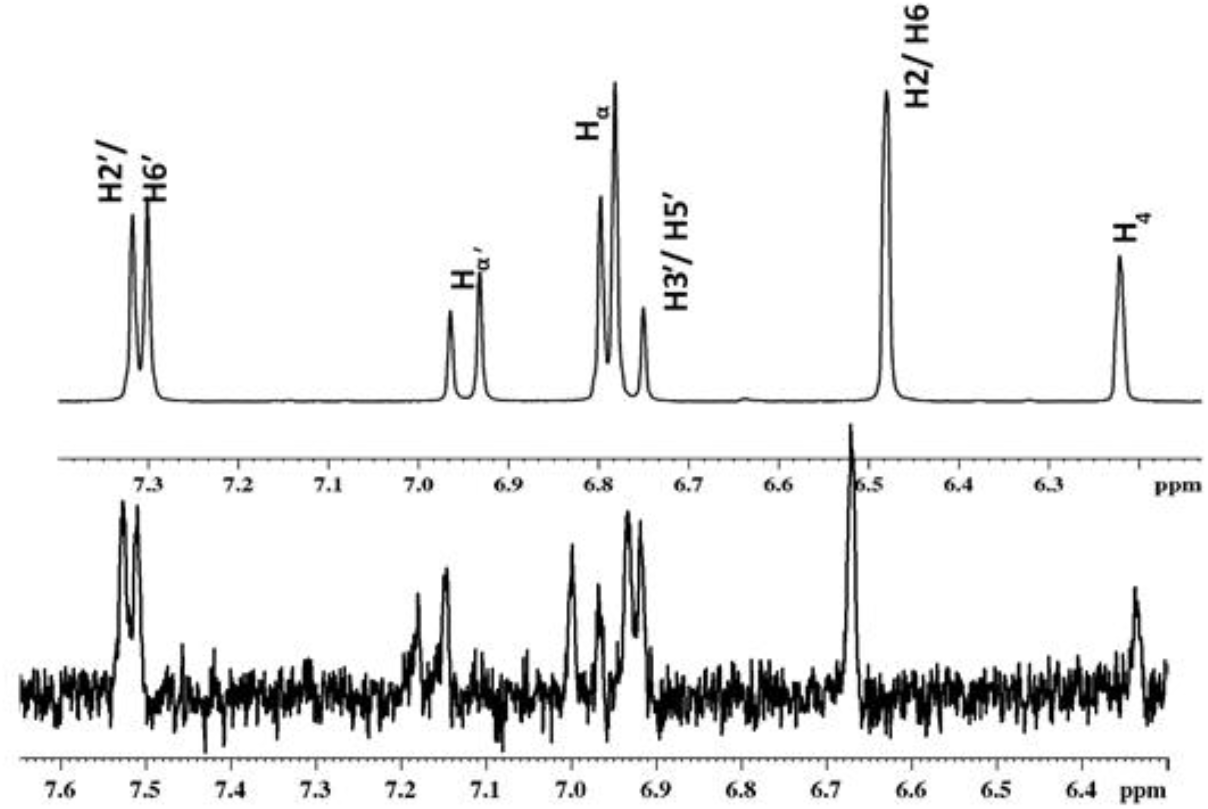
One dimensional proton spectrum of resveratrol in deuterated ethanol (above) and in D_2_O (below).

#### 3.5.3. DNA-resveratrol complex

Upon addition of resveratrol to d-(CCAATTGG)_2_ in small steps up to D/N ratio 1.0, a set of new signals corresponding to H2 and H6 protons of resveratrol appear (Fig.11a).

**Fig. 11.**
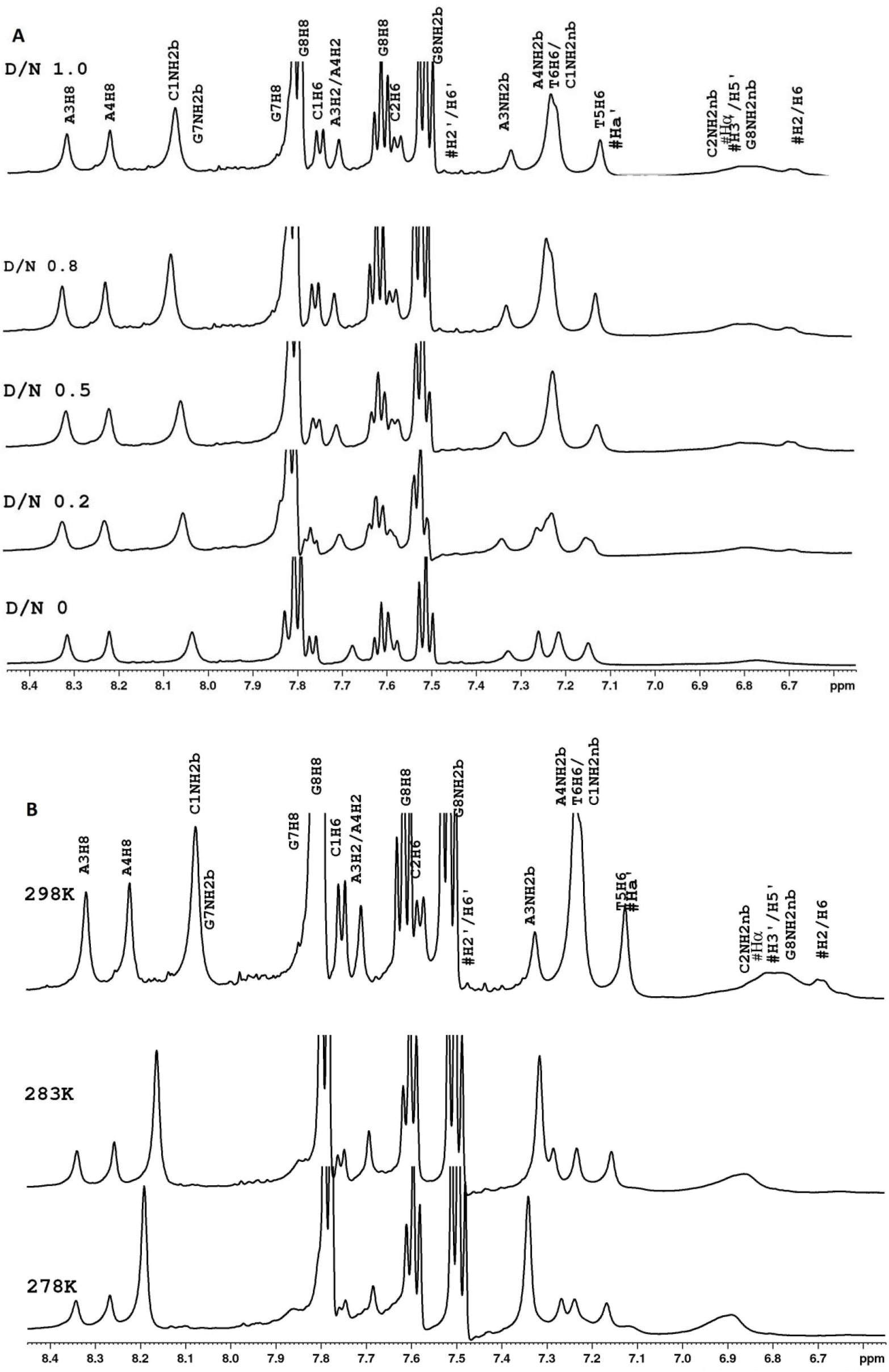
A) 1D 1H NMR spectrum of resveratrol-DNA complex at different D/N ratios, B) 1D 1H NMR spectrum of resveratrol-DNA complex at temperatures 278K, 283K and 298K. # denotes proton signals from resveratrol.

The change in chemical shift (Δδ) of d-(CCAATTGG)_2_ protons with increasing D/N ratio was gradual and small in magnitude (as shown in Supplimentary material table 1A and figure 11). T6H1 and T6H5” shifted upfield by ~0.04 and 0.06 ppm while A3H2 andA4H2 showed a downfield shift of ~ 0.03 ppm on binding to resveratrol.A4H6 was downfield shifted by 0.06 ppm.

All the spectral lines of DNA were somewhat uniformly broadened upon binding as the internal motions are affected, and protons get immobilized. Fig. 12 shows the broadening of spectral lines in the H1′ region. The very small chemical shift changes indicate that the interaction between resveratrol and d-(CCAATTGG)_2_ induced only a slight distortion in the DNA structure. ^1^H NMR spectra of the resveratrol-d-(CCAATTGG)_2_complex in the temperature range of 278-298 K (Fig.11b) show that drug protons are more prominent at 298K.

**Fig. 12.**
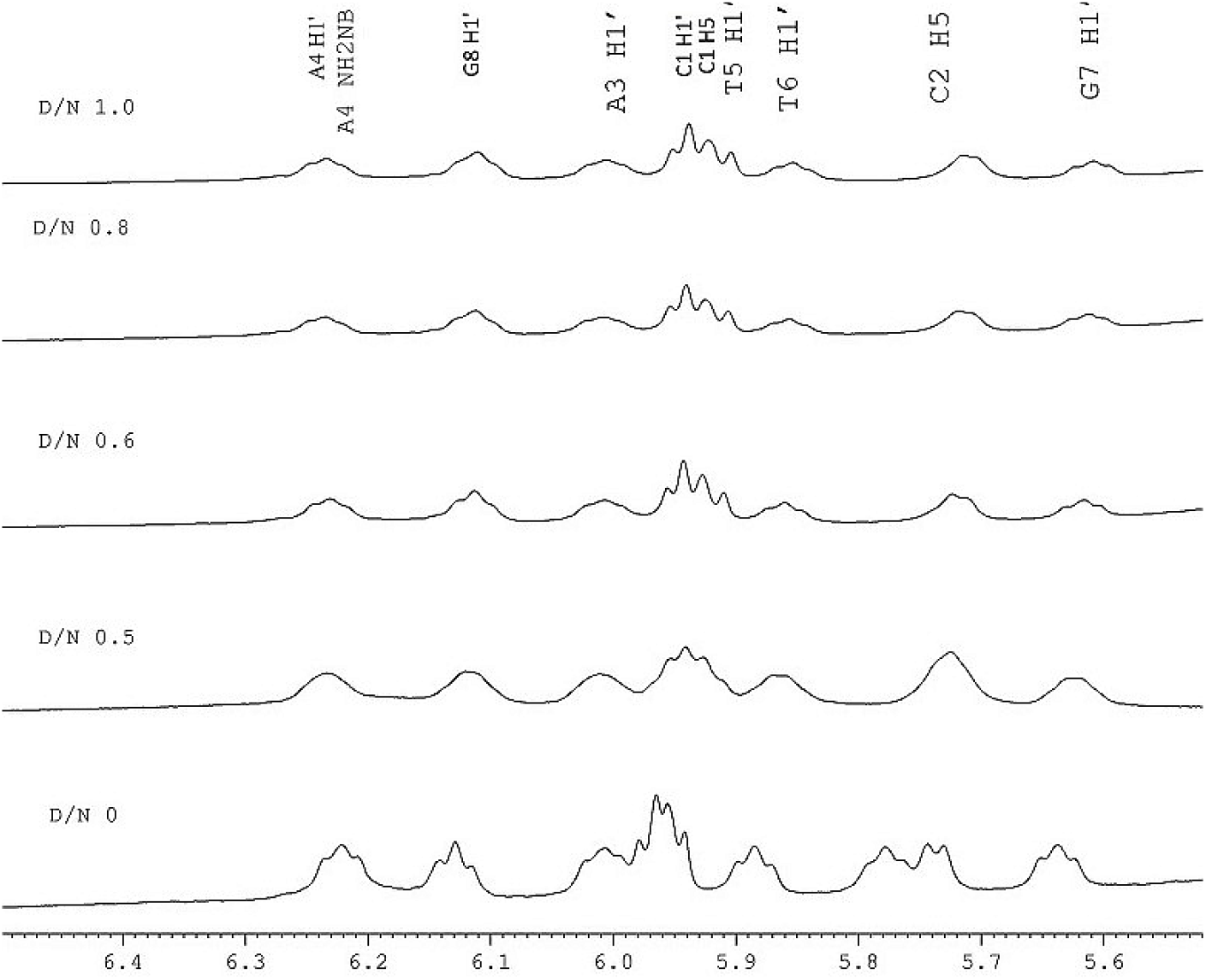
Sugar H1′ of DNA when complexed with resveratrol at different D/N ratios showing broadening of the peaks.

The 2D NOESY spectra of resveratrol d-(CCAATTGG)_2_ complex were obtained at different mixing times, D/N ratios and temperatures. Fig. 13 and Supplementary material Fig. 1A shows NOE connectivities observed in the resveratrol-d-(CCAATTGG)_2_ complex at D/N=1, 298 K andthe NOE peaks (all not shown) have been used in deriving the structure of the complex. The NOESY spectra were analyzed extensively at different mixing times. NOEs between pairs of proton G7NH^b^ - C2NH_2_^b^, C2NH_2_^nb^; T6NH - A3NH_2_^b^, A3NH_2_^nb^, A3H2; and T5NH-A4NH_2_^b^, A4NH_2_^nb^, A4H2 were observed establishing Watson-Crick base pairing between C2…. G7, T6….A3 and T5….A4 base pairs in the complex. This is furthersupported by the presence of intrastrand sequential connectivity T5NH-A4NH_2_^nb^ and interstrand sequential connectivities T6NH - C2NH_2_^b^, C2NH_2_^nb^ (Supplementary material Fig.1B). The sequential connectivities G7NH - T6NH, T6NH - T5NH were also observed. All the sequential connectivities between Base H8/H6/CH_3_/H5 and sugar protons H′, H2′, H2” were observed at all basepair steps. The NOE data therefore proves the existence of B-DNA duplex with no possibility of opening of base pairs. Hence, the intercalation mode of interaction between resveratrol and DNA in the ligand-DNA complex is completely ruled out.

**Fig. 13.**
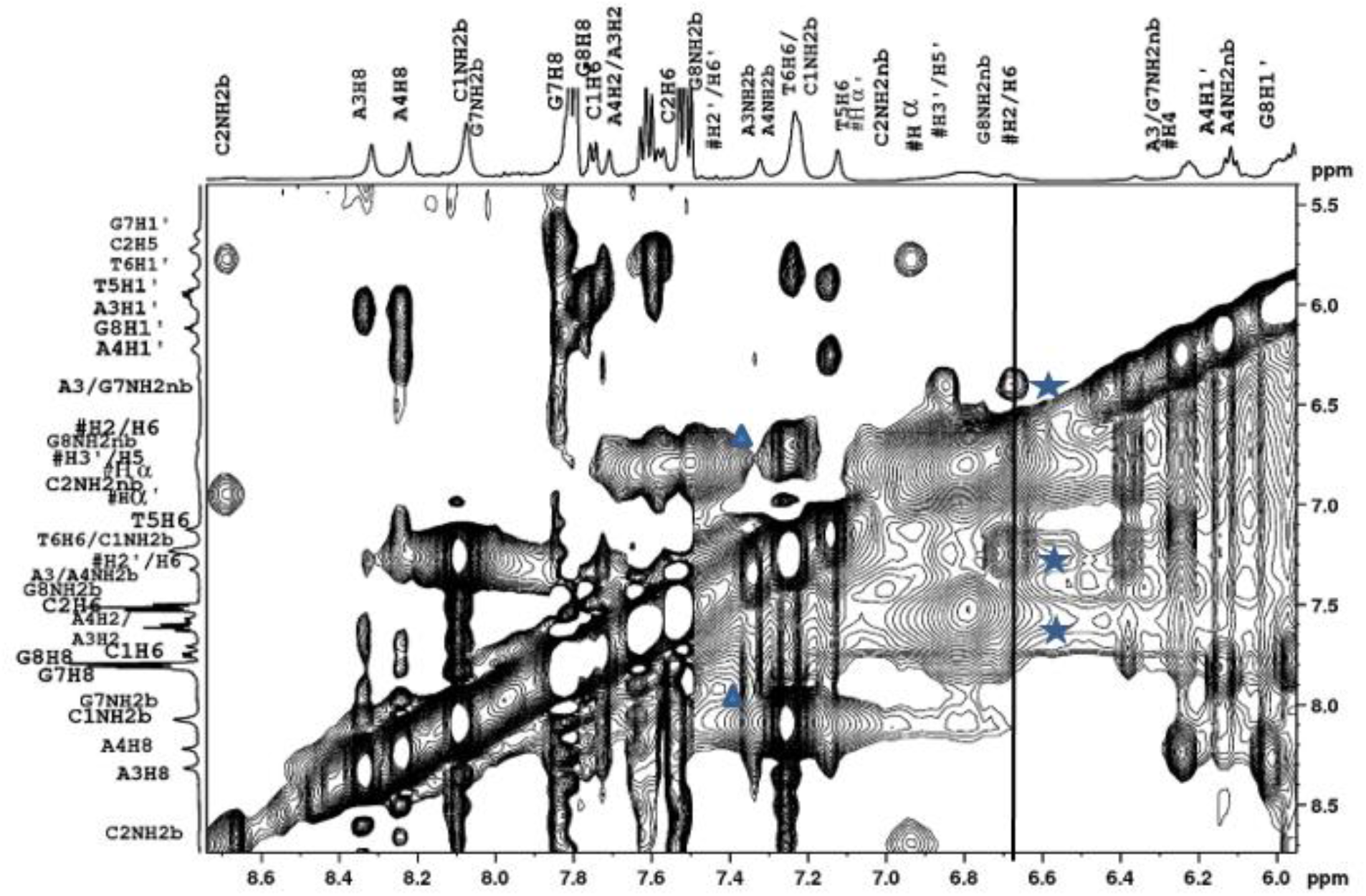
Expansion of the specific regions of the NOESY spectra of resveratrol complexed with d-(CCAATTGG)_2_ at D/N=1 at 298 K. 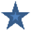 indicates intermolecular cross peaks between resveratrol and DNA protons, 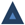 indicates intramolecular cross peaks of resveratrol.

Interaction between two molecules often causes changes in the chemical environment of the atoms at the interface, leading to chemical shift changes in proton NMR spectra. As a result, analysis of chemical shift can help one to pinpoint the sites of local interactions between the molecules. If the resonances that shift upon binding correspond to the residue directly involved in binding, chemical shift change can provide valuable insights into the structural basis for recognition. Hence can lead the way for designing or screening efforts focused on the discovery of novel ligands. In the present study, the slight shifts in G7H6, G7H2′, G7H1′, T6H6, T6H5”, A4H6, A4H2 and A3H2 protons suggests that binding affects the conformation of these bases of DNA. The shift in most of DNA protons is however insignificant, as has also been reported in earlier studies [69,70]. The distortions induced in DNA conformation appear to be relatively small in magnitude. The changes in resveratrol protons are not significant, ruling out the stacking mode of interaction. The NOE cross peaks were observed between the resveratrol H6/H2 protons and H6 of T6; NH_2_^nb^ of A3; H2 of A3 and A4 (overlapped) as indicated in figure 13. The small chemical shift changes are indicative of external binding of the resveratrol molecule near the AT base pair. The detailed structural information was, however, derived by analyzing NOEs. No evidence like intermolecular cross peaks (between resveratrol) was obtained to establish the binding of two molecules of resveratrol. Also, because of the self-complementary nature of the sequence, it can be judged that resveratrol forms cross peaks with NH_2_^nb^ of A3of first strand and H6 of T14 (i.e., T6 of the second strand). These intermolecular ^1^H-^1^H NOESY cross peaks (Supplementary material Table 1B) between resveratrol and DNA bases confirm that it binds near the AT base pair. Supplementary material Table 1C shows few intramolecular cross peaks observed for resveratrol in the complex spectrum.

### 3.6. Phosphorus-31 NMR Spectra

Phosphate resonances of uncomplexed DNA were assigned from the ^1^H-^31^P HMBC spectrum. C2pA3, G7pG8, C1pC2, T6pG7 and A3pA4 appear at −0.42, 0.52, −0.79, −0.85 and −1.01 ppm, respectively at 298 K while resonance of both A4pT5 and T5pT6 appear at −1.14 ppm. Upon binding of resveratrol to d-(CCAATTGG)_2_, the resonances broadened without any significant shift, as shown in Fig. 14. The largest shift (upfield) of 0.053 ppm was observed for T6pG7 followed by 0.017 ppm for A3pA4 while the A4pT5/T5pT6 resonance upfield shifted by 0.013 ppm (Supplementary material Table 3). No additional peaks were observed.

**Fig. 14.**
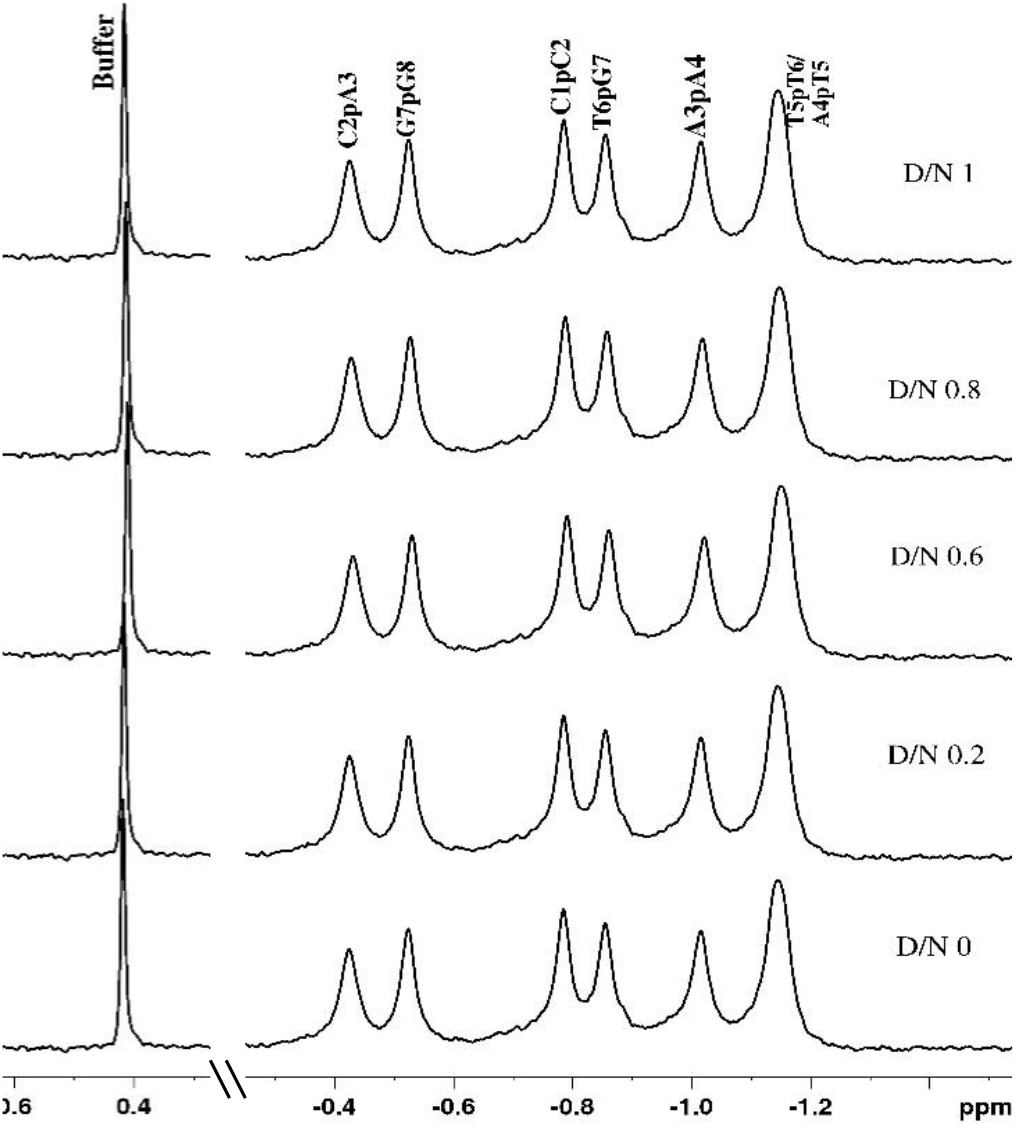
Proton decoupled ^31^P NMR spectra of resveratrol-(CCAATTGG)_2_complex as a function of added drug (D) to Nucleic acid duplex (N) stoichiometric ratios (D/N) of 0.0, 0.2, 0.4.0.6, 0.8 1.0 at 298 K in phosphate buffer.

It is known that ^31^P chemical shifts vary in response to local, sequence specific and induced environmental distortions in the duplex geometry[71],occur with ester O-P-O bond angle distortions or modifications in backbone torsional angles. The intercalating molecules induce deformation of phosphodiester chain, to allow the opening of base pairs, by converting from energetically more favorable BI conformation to a more flexible BII conformation. This usually results in a downfield shift more than1.5 ppm[72]. It is known that downfield shifts of ~1.5 ppm or more occur on the binding of DNA to classical intercalated[73,74], while ligands which bind externally to DNA such as pyridopurine[75], mitoxantrone[76],camptothecin[77], show much smaller shifts. Absence of any such large downfield shift in the present study excludes the possibility of opening of base pairs to allow intercalation. This result is in accord with the observation that the change in the absorption maximum on the binding of resveratrol to DNA sequence is insignificant as compared to that of 25-44 nm observed on the binding of typical intercalator to DNA[78,79]. Purely electrostatic associations between drug and nucleic acid, on the other hand, produce only small and generally upfield^31^P chemical shifts [80]which are not observed in the present study. Thus the association of resveratrol with octamer does not involve direct intercalation or electrostatic interaction. The minute chemical shiftsfound thus corroboratefor the external binding of the compound. The small shifts in the positions of T6pG7, A3pA4 and A4pT5/T5pT6 indicate that resveratrol is binding near the AATT region.

### 3.7. Observations from NMR and other spectral data

Analysis of the shifts in proton NMR structure indicates no substantial changes in the DNA conformation upon binding with resveratrol. Chemical shift changes in the base protons of T5, T6, G7, A3 and A2, though small, bespoke the binding of the compound near these bases. The NOE cross peaks obtained with the resveratrol H2/H6 protons substantiate these findings and confirms that resveratrol binding to the AATT minor groove region. The features of observed ^31^P resonances also vouch for the external binding of the compound. The absorption, fluorescence and CD spectral data described an external binding of the compound to the DNA sequence. These findings support our earlier findings of the external binding of resveratrol to two DNA sequences with central AATT segment [54].

### 3.8. Structure-Conformation and Dynamics

The sequenced-(CCAATTGG)_2_ was prepared from PDB data 1JTL. The terminal residues and ligand from the structure were deleted and used for further analysis. To obtain a complex structure, d-(CCAATTGG)_2_ was first docked with resveratrol. The docking conformations showed three clusters. The lowest energy structure from the most populated clusters was taken as the starting structure of the complex. The complex structure from docking was curated for DNA and resveratrol structure using ambpdb and antechamber protocol for amber compatible structure. Modified starting structure of the complex was prepared. The structure was energy minimized and MD simulations were performed.

#### 3.8.2. Molecular Dynamics (MD) simulations

MD simulations were carried out using the sander module in AMBER18 simulation package. Hydrogen atoms were added to both ligand and DNA. The leaprc force fields, general amber force field (GAFF) and bsc1DNA were used for ligand and DNA, respectively in the tleap module[81]. The complex structure was solvated in a truncated octahedron periodic cell centered on DNA and explicit water molecules were added using the TIP3P water model [82]. tleap has added around 5710water molecules within 15Å of solvation between the solute (DNA) and the edges of the box. The solute was neutralized with 14 sodium ions making the box size 72.86 × 72.86 × 72.86 Å. Minimization and equilibration of the DNA-resveratrol complex were carried out followed by MD production run. Firstly, the DNA and the ligand were minimized with both the molecules restrained for 1000 steps, with a cut off of 10 Å. The minimized structure was subjected to another round of minimization without any restraint either on DNA or on resveratrol. This is followed by heating the system using the NVT ensemble and the SHAKE algorithm to constrain hydrogen bonds[83]. The temperature was increased from 100K to 300K during the first 10 ps (5000 steps) and kept constant at 300K for the next 90 ps. The system was then equilibrated after reducing the constraint to 0.5 kcal/mol in the NPT ensemble at constant temperature (300K) and pressure (1 atm) for 50ps. The production phase of the simulation was performed in the NVT ensemble at 300 K and Langevinthermostat with collision frequency 1ps^−1^. Distance restraints obtained from NOE intensities were introduced using make DIST_RSTprogram in Amber and simulation was carried out for 1ns with a time step of 2 fs. A total of 223 distances were used. Root mean square deviation and backbone torsional angles were estimated using the cpptraj module [84].

The 1 ns trajectory was stable, as indicated by the steady root mean square deviation (rmsd) values, as shown in Fig. 15. The rmsd was between 2.5 Å and 3.5 Å considering all 8 residues and 2 Å to 2.5Å for central residues (red line). The potential and kinetic energies of the system were stabilized after the initial 20 ps equilibration run. The minimum energy structures were generated using cpptraj, and the root mean square deviation (rmsd) of nine lower energy structures with the lowest energy structure was obtained as 1.709 Åandthe superimposed minimum energy structures are shown in Fig. 15C. Pseudo puckering of atom C1′ through O4′for residues 3 to 6 is shown in Fig. 15B. Sugar puckering of residues 3, 5 and 6 fall in the range of 150 degrees while residue 4 shows a slightly higher value as displayed in the Fig. 15B. This could be due to the binding of resveratrol near this residue. These values are consistent with B DNA parameter. The lowest energy structure is stabilized with a total potential energy = −57746.5 kcal/mol. In the lowest energy structure, resveratrol 3OH forms four hydrogen bonds with DNA namely, O with G15H6 and A3NH2; while H with O4′ of G15sugar and O5′of T14sugar. All distances are within 2.7 Å. This reveals that resveratrol is accommodated to the AT region with one terminal fixed by making hydrogen bonds, thereby stabilizing the structure. The chemical shift changes observed for T6 and A3 base and sugar protons as well as phosphorus also support this. The binding of resveratrol near these bases induce the changes, though the shifts are very small. Increase in the fluorescence intensity of resveratrol upon binding, as described earlier, is because of the intrusion of the OH group into the minor groove region.

**Fig. 15.**
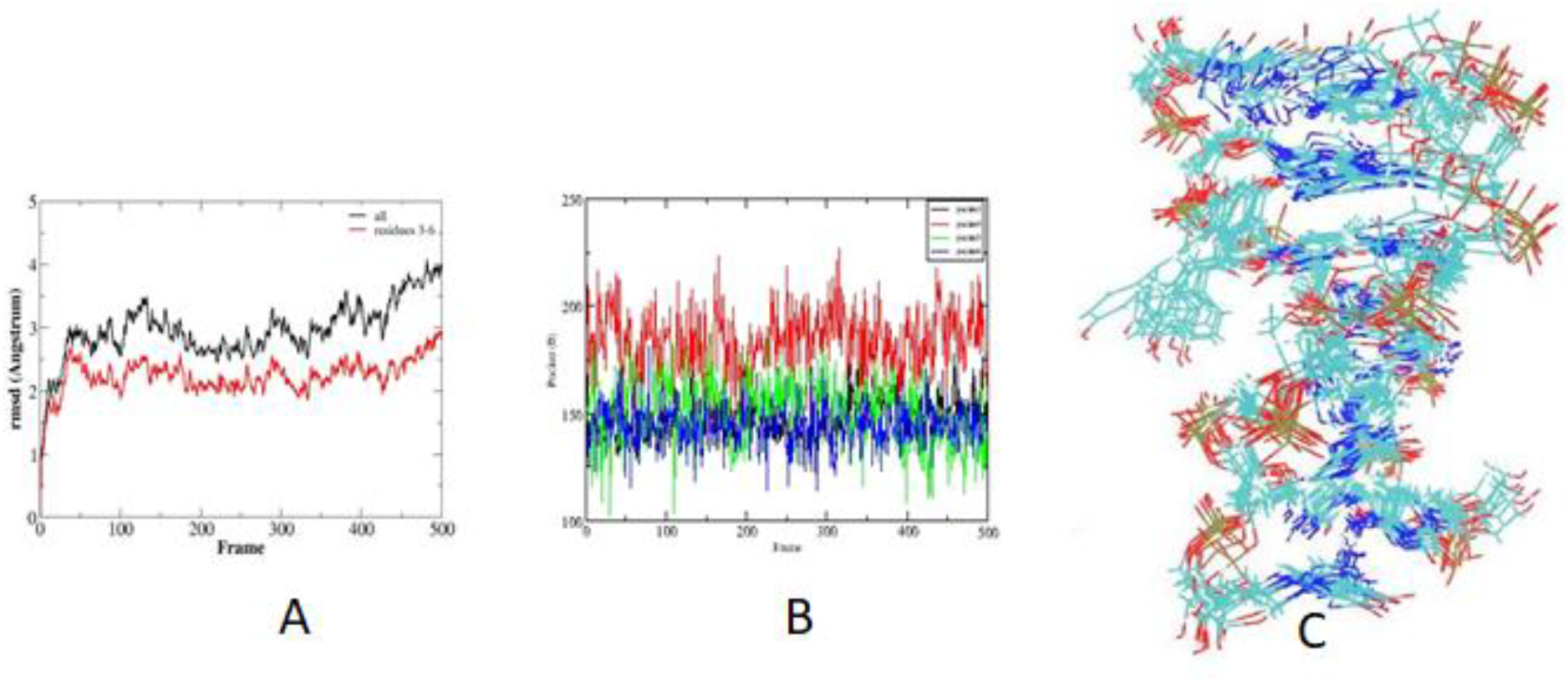
A) rmsd when all residues are considered (black) and when only central four residues are considered (red), B) sugar puckering of residues 3, 4, 5 and 6 and C) nine lowest energy structures obtained after MD simulation superimposed.

Structural parameters of DNA were analyzed using Curves+ software. The helical analysis shows an average twist of 20.2° and helical rise of 3.23 Å. All sugars except the ones for A4 on the first strand and T13 on the second strand are in C2′endo puckering, retaining the features of B-DNA. Minor groove width of DNA was increased as shown in Fig. 16. Minor groove depth increased slightly. Major groove width of AT base pair slightly increased while the depth decreased. This might have caused because of the binding of resveratrol in the minor groove. Similar features were indicated in the literature, in the case of minor groove binding agents [85–88].

**Fig. 16.**
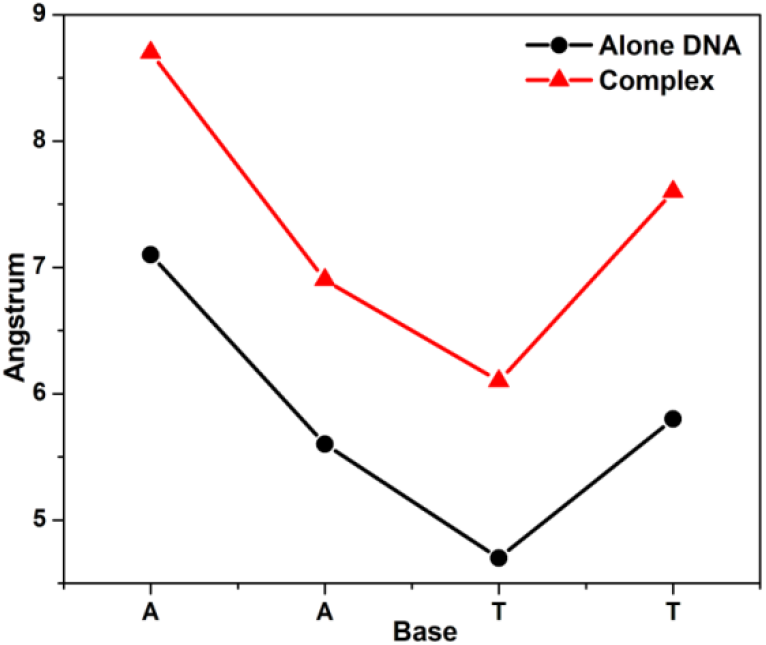
Changes in the minor groove width of DNA alone and when complexed with resveratrol.

## 4. Conclusion

The paper discusses the interaction of resveratrol with an octamer sequence which is present in the promoter region of various oncogenes and many nuclear factors. Absorption and fluorescence studies revealed that the compound is interacting externally. Fluorescence technique which is more sensitive than absorption shows that resveratrol binds with moderate to high affinity to the octamer sequence. Also, the high fluorescence exhibited by resveratrol in the presence of DNA can be used as a practical DNA light probe. Circular dichroism data display moderate changes in the DNA conformation upon binding. The small changes in the melting temperatures of DNA and its 1:1 and 1:2 complexes with resveratrol upholds the spectroscopic observations that resveratrol binds externally to DNA. NOE cross peak between H2/H6 protons of resveratrol with A3, A4 and T6 base protons establishes the formation of resveratrol-DNA complex at the minor groove. Subtle changes in the chemical shifts corroborate for the binding. Lowest energy structure obtained from the molecular dynamics analysis confirms that the DNA conformation changed subtly to accommodate the compound, supports with the findings of circular dichroism data. The complex structure was stabilized by hydrogen bonds formed between one terminal end of resveratrol with DNA. These results endorse resveratrol as a minor groove binding agent and pave the way to develop resveratrol and its analogues as suitable candidates as a new class of DNA sensing probes. Many minor groove binders are known for their antibacterial activity. The results obtained urge one to probe the effect resveratrol binding to specific nucleotide sequences which play an essential role in gene regulation and potential candidate as a novel antibacterial/anticancer drug.

## Acknowledgement

SK acknowledges the University GrantCommission (UGC) for a student research fellowship.MN appreciatesthe Council of Scientific and Research Council (CSIR), India for partial funding. MN is thankful to Dr David A. Case, for providing a free license of Amber18 package.

## Authorship contribution

PK performed NMR experiments and analyzed data, SK performed absorption, fluorescence, CD, and DSC experiments. MN conceptualized, designed and analyzed the data and wrote the manuscript.

## Supplementary material

**Supplementary material Table 1A.**
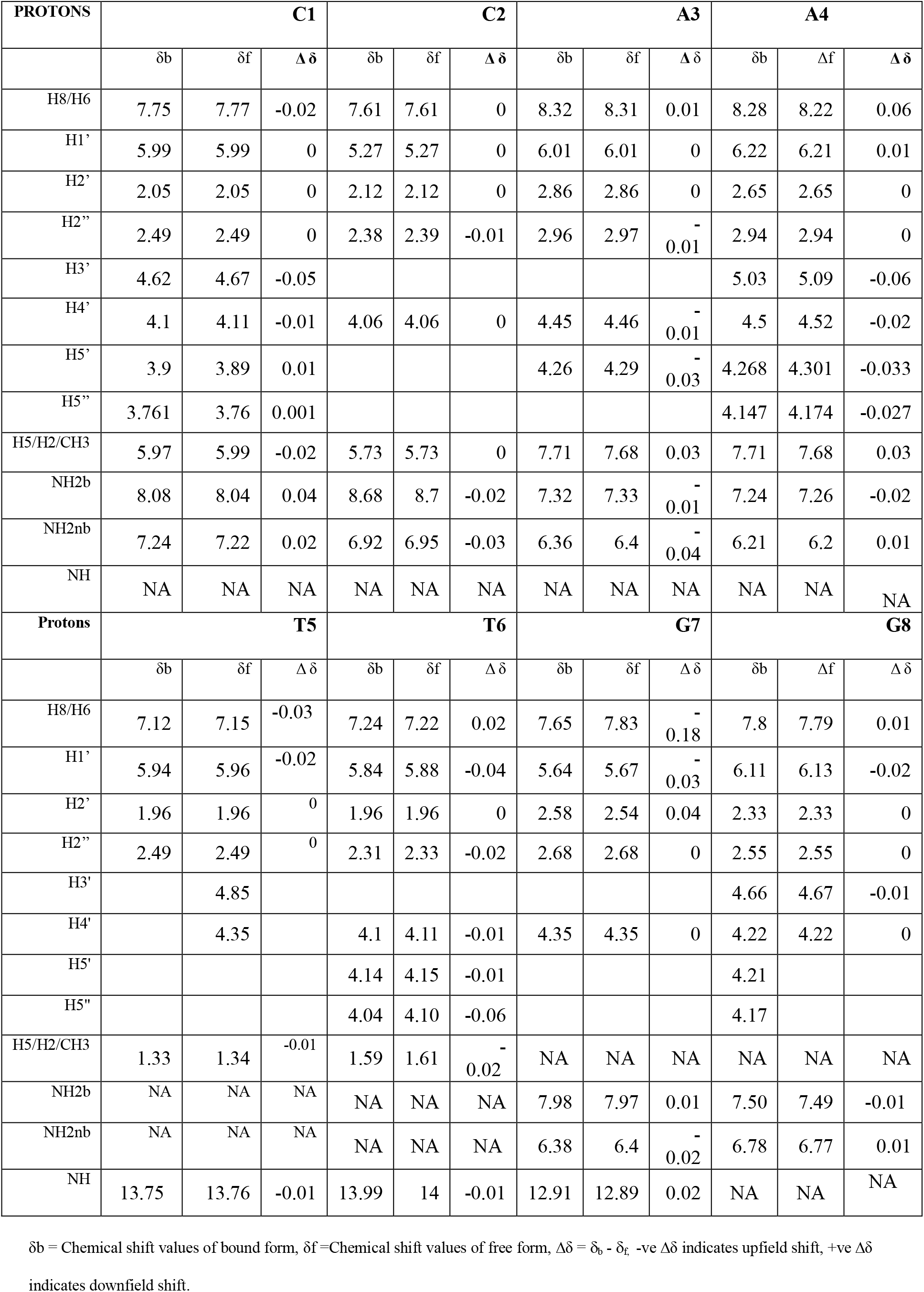
Chemical shift values (ppm) of d-(CCAATTGG)_2_ in 90%H_2_0 +10%D_2_0 at 298K.

**Supplementary material Table 1B.**
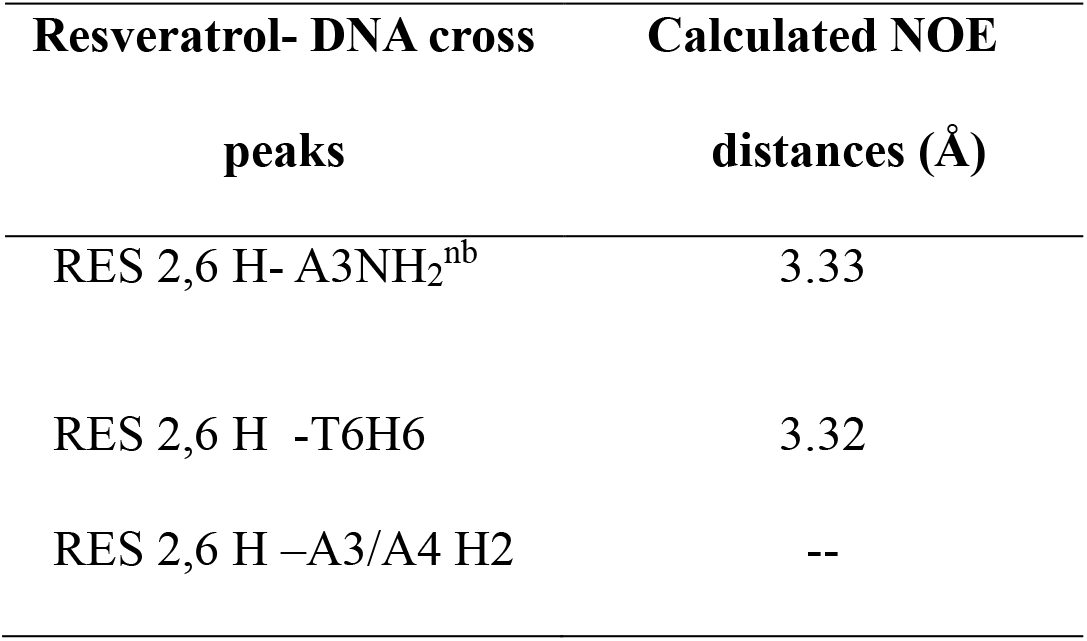
Intermolecular cross peaks between resveratrol and d-(CCAATTGG)_2_.

| **Resveratrol- DNA cross** | **Calculated NOE** |
| --- | --- |
| **peaks** | **distances (Å)** |
| RES 2,6 H- A3NH<sub>2</sub><sup>nb</sup> | 3.33 |
| RES 2,6 H -T6H6 | 3.32 |
| RES 2,6 H –A3/A4 H2 | -- |

**Supplementary material Table 1C.**
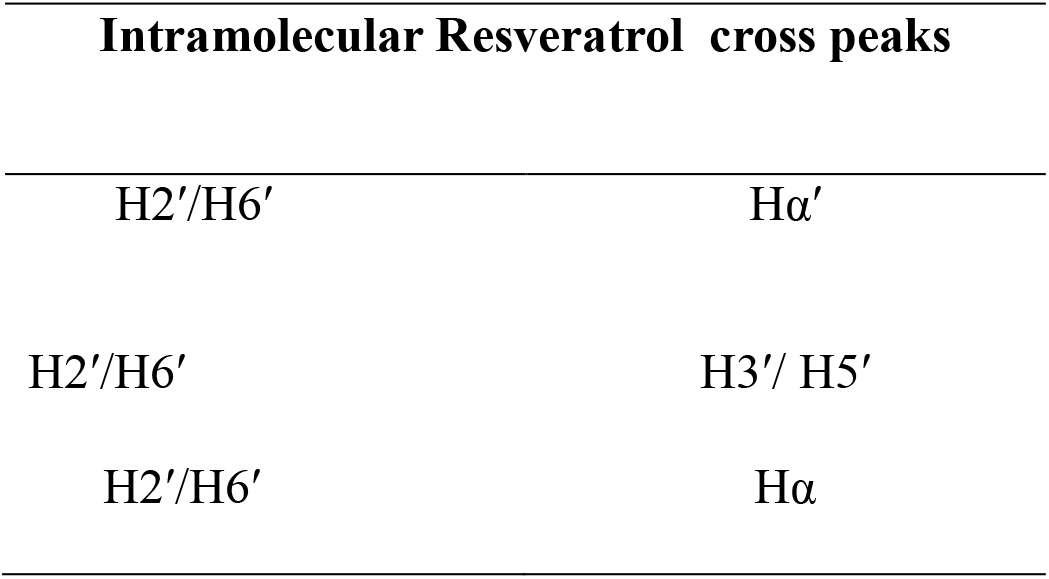
Intramolecular cross peaks within resveratrol, observed in complex spectrum.

| Intramolecular Resveratrol cross peaks |  |
| --- | --- |
| H2'/H6' | Hα' |
| H2'/H6' | H3'/ H5' |
| H2'/H6' | Hα |

**Supplementary material Table 2.**
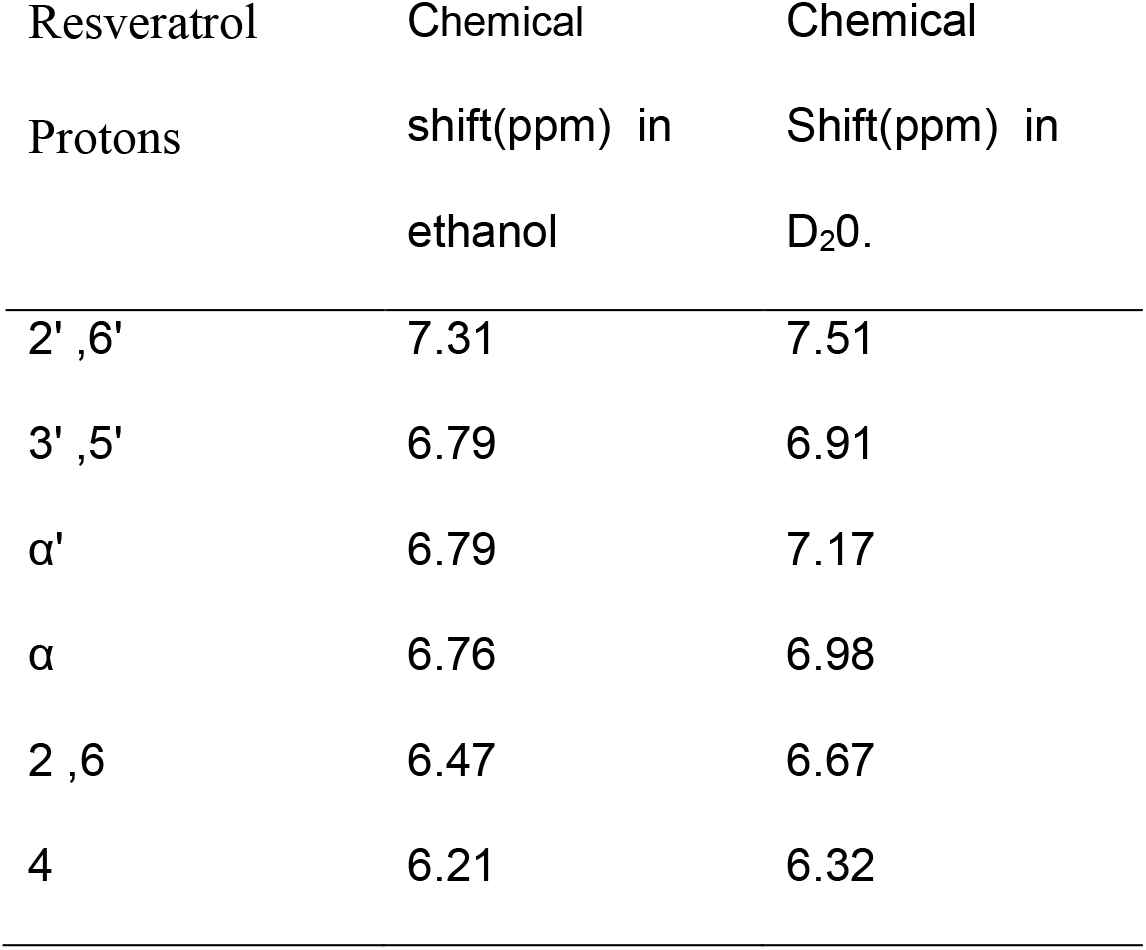
Chemical Shifts of resveratrol protons in deuterated ethanol and D20.

**Supplementary material Table 3.**
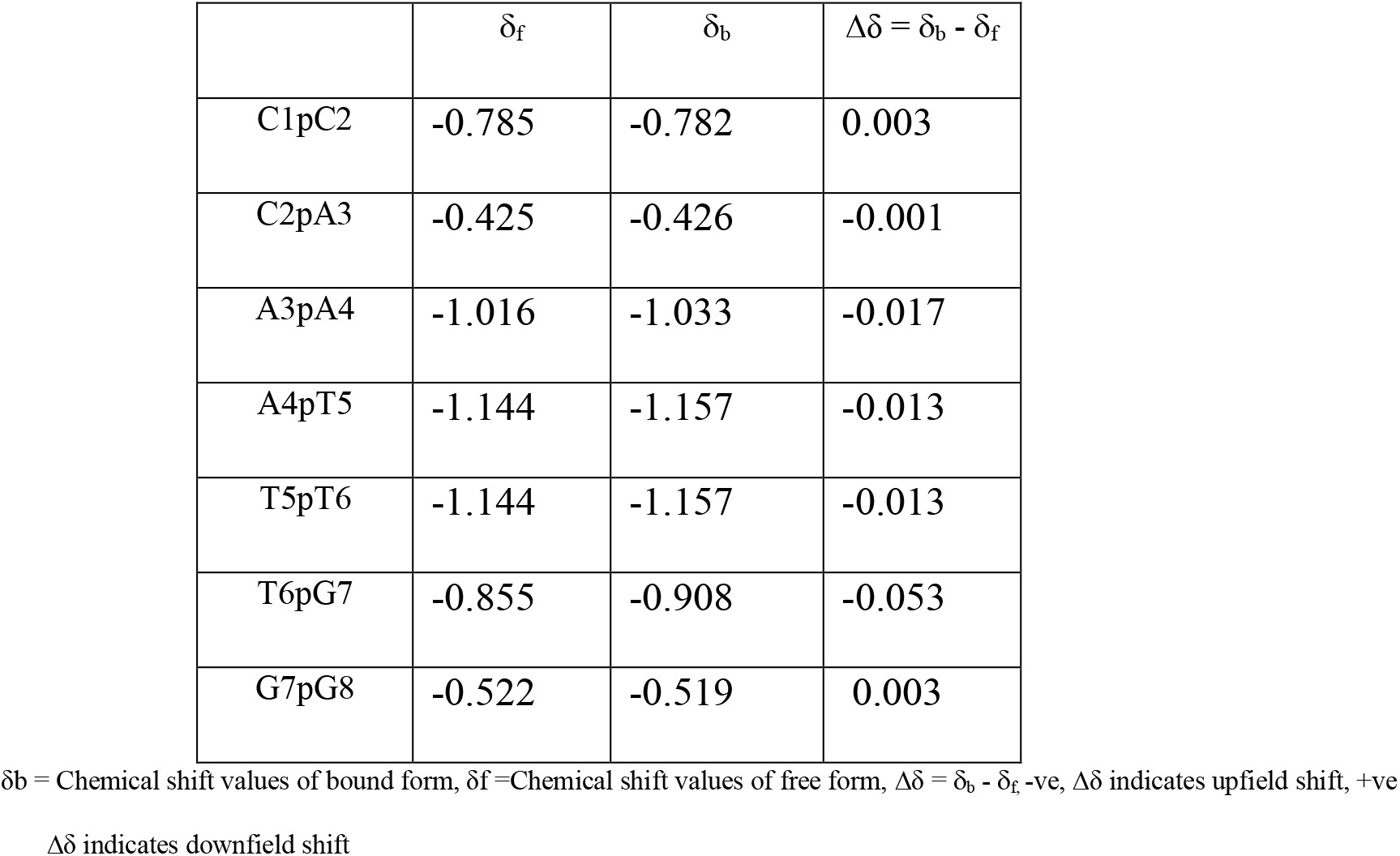
Chemical shift of^31^P resonances (ppm) of DNA octamer in d-(CCAATTGG)_2_complexed with resveratrol at 298K

**Supplementary material Fig.1A.**
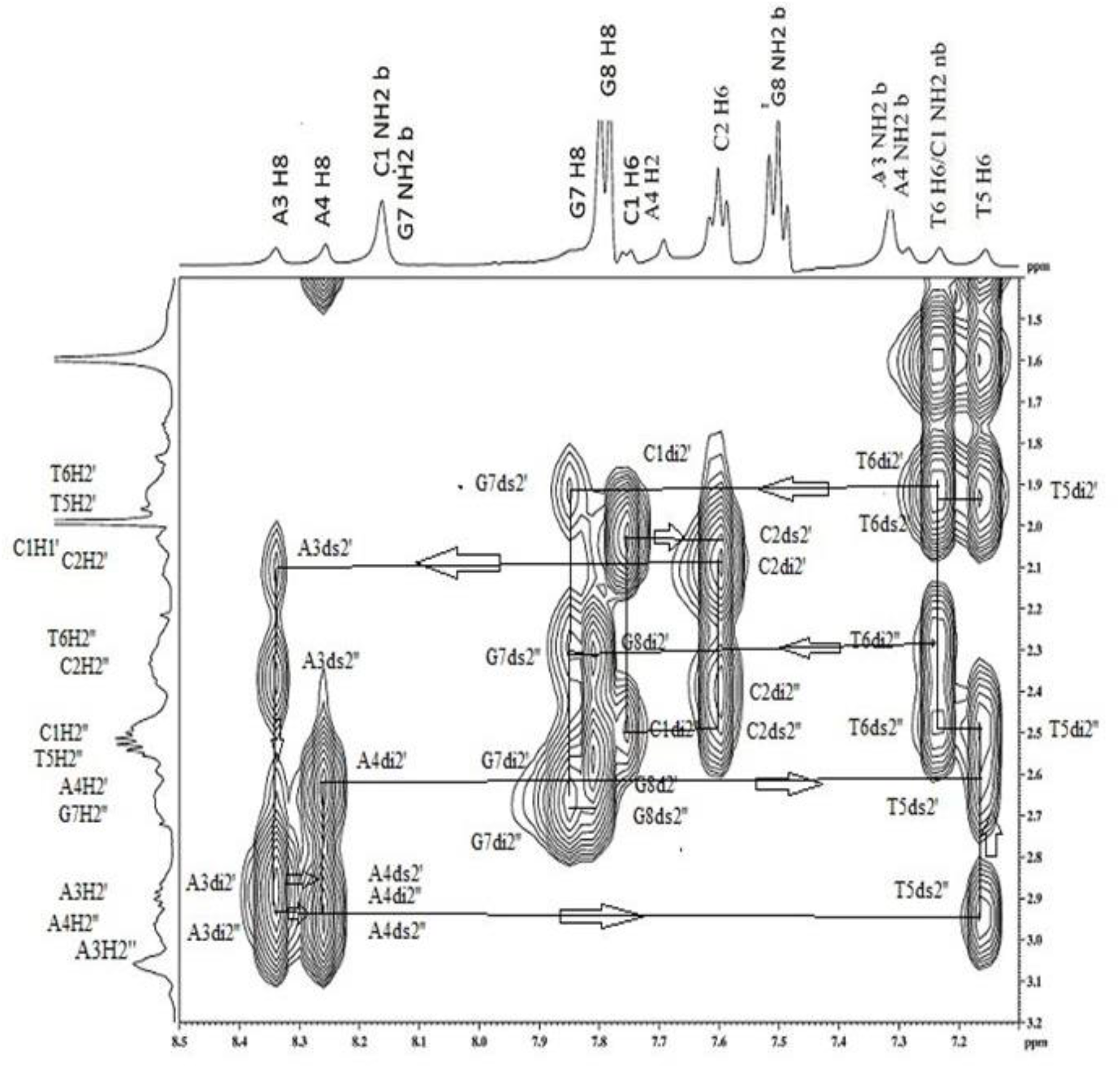
NOE connectivities of resveratrol-DNA complex showing base-H2′-H2′ region.

**Supplementary material Fig.1B.**
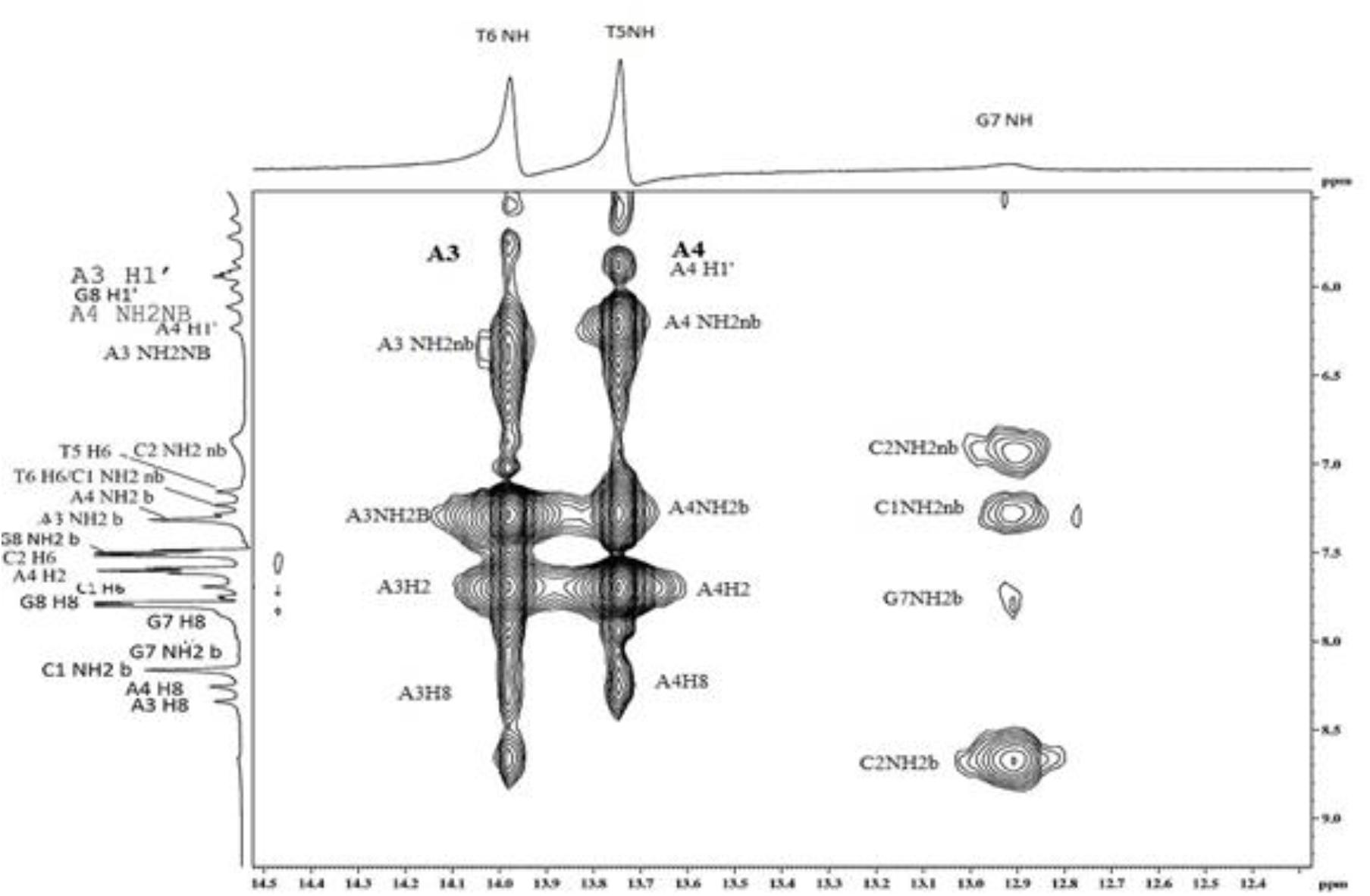
NOE connectivities of resveratrol-DNA complex showing base-NH region.

## References

[1] D. Gibson, Drug-DNA interactions and novel drug design, Pharmacogenomics J. vol. 2 (2002) 275–276, doi: 10.1038/sj.tpj.6500133.

[2] O. Kennard, DNA-drug interactions, Pure Appl. Chem. 65 (1993) 1213–1222, doi: https://doi.org/10.1351/pac199365061213.

[3] R. Palchaudhuri, P. J. Hergenrother, DNA as a target for anticancer compounds: methods to determine the mode of binding and the mechanism of action, Current Opinion in Biotechnology. 18 (2007) 497–503, doi: 10.1016/j.copbio.2007.09.006.

[4] T. Sarwar, S. U. Rehman, M. A. Husain, H. M. Ishqi, M. Tabish, Interaction of coumarin with calf thymus DNA: Deciphering the mode of binding by in vitro studies, Int. J. Biol. Macromol. 73 (2015) 9–16, doi: 10.1016/j.ijbiomac.2014.10.017.

[5] J. Ren, J. B. Chaires, Sequence and structural selectivity of nucleic acid binding ligands, Biochemistry. 38 (1999) 16067–16075, doi: 10.1021/bi992070s.

[6] G. H. Kocharyan, S. H. Minasyan, L. A. Tavadyan, Interaction of flavonoids: morin, quercetin and rutin, with DNA, Chemistry and Biology. 1 (2016) 49–54, corpus ID: 461125.

[7] J. Gusman, A reappraisal of the potential chemopreventive and chemotherapeutic properties of resveratrol, Carcinogenesis. 22 (2001) 1111–1117, doi: 10.1093/carcin/22.8.1111.

[8] C. N. N’Soukpoé-Kossi, P. Bourassa, J. S. Mandeville, L. Bekale, H. A. Tajmir-Riahi, Structural modeling for DNA binding to antioxidants resveratrol, genistein and curcumin, J. Photochem. Photobiol. B Biol. 15 (2015) 69–75, doi: 10.1016/j.jphotobiol.2015.07.007.

[9] H. Xiang, Y. Xie, J. Huang, Study on the interaction of DNA with resveratrol by resonance light scattering technique and its analytical application, J. Anal. Chem. 66 (2011) 618–622, doi: 10.1134/S1061934811040071.

[10] B. C. Trela, A. L. Waterhouse, Resveratrol: Isomeric Molar Absorptivities and Stability, J. Agric. Food Chem. 44 (1996) 1253–1257, https://pubs.acs.org/sharingguidelines.

[11] J. Gambini, M. Ingles, G. Olaso, R. Lopez-Grueso, V. Bonet-Costa, L. Gimeno-Mallench, C. Mas-Bargues, K. M. Abdelaziz, M. C. Gomez-Cabrera, J. Vina, C. Borras, Properties of Resveratrol: In Vitro and In Vivo Studies about Metabolism, Bioavailability, and Biological Effects in Animal Models and Humans, Oxidative Medicine and Cellular Longevity. 2015 (2015) 13, doi: 10.1155/2015/837042.

[12] X. Zheng, W. D. Meng, Y. Y. Xu, J. G. Cao, F. L. Qing, Synthesis and anticancer effect of chrysin derivatives, Bioorganic Med. Chem. Lett. 13 (2003) 881–884, doi: 10.1016/S0960-894X(02)01081-8.

[13] S. Leone, T. Cornetta, E. Basso, R. Cozzi, Resveratrol induces DNA double-strand breaks through human topoisomerase II interaction, Cancer Lett. 295 (2010) 167–172, doi: 10.1016/j.canlet.2010.02.022.

[14] S. K. Singh, S. Banerjee, E. P. Acosta, J. W. Lillard, R. Singh, Resveratrol induces cell cycle arrest and apoptosis with docetaxel in prostate cancer cells via a p53/ p21 WAF1/CIP1 and p27 KIP1 pathway, Oncotarget. 8 (2017) 17216–17228, www.impactjournals.com/oncotarget.

[15] A. liontas, H. yeger, Curcumin and resveratrol induce apoptosis and nuclear translocation and activation of p53 in human neuroblastoma, Anticancer Research. 24 (2004) 987–998.

[16] K. H. Kang, H. Kim, K. H. Choi, p21 Promotes Ceramide-Induced Apoptosis and Antagonizes the Antideath Effect of Bcl-2 in Human Hepatocarcinoma Cells, Experimental Cell Research. 253 (1999) 403–412, http://www.idealibrary.com.

[17] M. T. Laux, M. Aregullin, J. P. Berry, J. A. Flanders, E. Rodriguez, Identification of a p53-Dependent Pathway in the Induction of Apoptosis of Human Breast Cancer Cells by the Natural Product, Resveratrol, J. Altern. Complement. Med. 10 (2004) 235–239, doi: 10.1089/107555304323062211.

[18] M. Jang, L. Cai, G. O. Udeani, K. V. Slowing, C. F. Thomas, C. W. Beecher, H. H. Fong, N. R. Farnsworth, A. D. Kinghorn, R. G. Mehta, R. C. Moon, J. M. Pezzuto, Cancer chemopreventive activity of resveratrol, a natural product derived from grapes, Science. 275 (1997) 218–220, doi: 10.1126/science.275.5297.218.

[19] S. Usha, I. M. Johnson, R. Malathi, Modulation of DNA intercalation by resveratrol and genistein, Mol. Cell. Biochem. 284 (2006) 57–64, doi: 10.1007/s11010-005-9013-6.

[20] S. Usha, I. M. Johnson, R. Malathi, Interaction of resveratrol and genistein with nucleic acids, J. Biochem. Mol. Biol. 38 (2005) 198–205, doi: 10.5483/bmbrep.2005.38.2.198.

[21] P. Bucher, Weight matrix descriptions of four eukaryotic RNA polymerase II promoter elements derived from 502 unrelated promoter sequences, J. Mol. Biol. 212 (1990) 563–578, doi: 10.1016/0022-2836(90)90223-9.

[22] A. Efstratiadis, J. W. Posakony, T. Maniatis, R. M. Lawn, C. Connell, R. A. Spritz, J. K. Deriel, B. G. Forget, S. M. Weissman, J. L. Slightom, A. E. Blechl, O. Smithies, F. E. Baralle, C. C. Shoulders, N. J. Proudfoot, The structure and evolution of the human β-globin gene family, Cell. 21 (1980) 653–668, doi: 10.1016/0092-8674(80)90429-8.

[23] R. Mantovani, A survey of 178 NF-Y binding CCAAT boxes, Nucleic Acids Res. 26 (1998) 1135–1143, doi: 10.1093/nar/26.5.1135.

[24] S. Borghini, M. Vargiolu, M. D. Duca, R. Ravazzolo, I. Ceccherini, Nuclear factor Y drives basal transcription of the human TLX3, a gene overexpressed in T-Cell Acute Lymphocytic Leukemia. Mol. Cancer. Res. 9 (2006) 635–643,doi: 10.1158/1541-7786.MCR-05-0250.

[25] P. P. Kuang, M. Joyce-Brady, X. H. Zhang, J. C. Jean, R. H. Goldstein, Fibulin-5 gene expression in human lung fibroblasts is regulated by TGF-β and phosphatidylinositol 3-kinase activity, Am. J. Physiol. -Cell Physiol. 291 (2006) 1412–1421, doi: 10.1152/ajpcell.00087.2006.

[26] S. Abhiman, L. M. Iyer, L. Aravind, A. Bateman, BEN: a novel domain in chromatin factors and DNA viral proteins, Bioinformatics. 24 (2008) 458–461, doi: 10.1093/bioinformatics/btn007.

[27] Q. Dai, A. Ren, J. O. Westholm, A. A. Serganov, D. J. Patel, E. C. Lai, The BEN domain is a novel sequence-specific DNA-binding domain conserved in neural transcriptional repressors, Genes and Dev. 27 (2013) 602–614, doi: 10.1101/gad.213314.113.

[28] C. Xuan, Q. Wang, X. Han, Y. Duan, L. Li, L. Shi, RBB, a novel transcription repressor, represses the transcription of HDM2 oncogene, Oncogene. 32 (2013) 3711–3721, doi: 10.1038/onc.2012.386.

[29] Q. Dai, Q, A. Ren, A, J.O. Westholm, H. Duan, D. J. Patel, E.C. Lai, Common and distinct DNA-binding and regulatory activities of the BEN-solo transcription factor family, Genes and Development. 29 (2015) 48–62, doi: 10.1101/gad.252122.114.

[30] S. Ratner, K. Murakami-Murofushi, A new radiochemical assay for argininosuccinase with purified [^14^C]argininosuccinate, Anal. Biochem. 106 (1980) 134–147, doi: 10.1016/0003-2697(80)90129-3.

[31] T. Matsubasa, M. Takiguchi, I. Matsuda, M. Mori, Rat ArgininosuccinateLyase Promoter: The Dyad-Symmetric CCAAT Box Sequence CCAATTGG in the Promoter Is Recognized by NF-Y, J.Biochem. 116 (1994) 1044–1055, https://academic.oup.com/jb/article-abstract/116/5/1044/941364.

[32] J. B. Park, M. Levine, Characterization of the promoter of the human ribonucleotide reductase R2 gene, Biochem. Biophys. Res. Commun. 267 (2000) 651–657, doi: 10.1006/bbrc.1999.1980.

[33] P. Borger, H. Matsumoto, S. Boustany, M.M.C. Gencay, J. K. Burgess, G. G. King, J. L. Black, M. Tamm, M. Roth, Disease-specific expression and regulation of CCAAT/enhancer-binding proteins in asthma and chronic obstructive pulmonary disease, J. Allergy Clin. Immunol. 119 (2007) 98–105, doi: 10.1016/j.jaci.2006.07.056.

[34] I. Panagopoulos, F. Mertens, Characterization of the human CREB3L2 gene promoter, Oncol. Rep. 21 (2009) 615–624, doi: 10.3892/or_00000264.

[35] C. Santoro, N. Mermod, P. C. Andrew, R. Tjian, A family of human CCAAT-box-binding proteins active in transcription and DNA replication: cloning and expression of multiple cDNAs, Nature. 334 (1988) 218–224, doi: 10.1038/334218a0.

[36] K. Uytterhoeven, J. Sponer, L. V. Meervelt, Two 1:1 binding modes for distamycin in the minor groove of d(GGCCAATTGG), Eur. J. Biochem. 269 (2002) 2868–2877, doi: 10.1046/j.1432-1033.2002.02952.x.

[37] M. Coll, J. Aymami, G. A. V. Marel, J. H. V. Boom, A. Rich, A. H. J. Wang, Molecular Structure of the Netropsin-d(CGCGATATCGCG)2 Complex: DNA Conformation in an Alternating AT Segment, Biochemistry. 28 (1989) 310–320, doi: 10.1021/bi00427a042.

[38] K. V. Hecke, P. C. Nam, M. T. Nguyen, L. V. Meervelt, Netropsin interactions in the minor groove of d(GGCCAATTGG)2 studied by a combination of resolution enhancement and ab initio calculations, FEBS J. 272 (2005) 3531–3541, doi: 10.1111/j.1742-4658.2005.04773.x.

[39] M. S. Searle, K. J. Embrey, Sequence-specific interaction of Hoescht 33258 with the minor groove of an adenine-tract DNA duplex studied in solution by 1 H NMR spectroscopy, Nucleic Acids Research. 18 (1990) 3753–3762, doi.org/10.1093/nar/18.13.3753.

[40] P. E. Pjura, K. Grzeskowiak, R. E. Dickerson, Binding of Hoechst 33258 to the minor groove of B-DNA, J. Mol. Biol. 197 (1987) 257–271, doi: 10.1016/0022-2836(87)901239.

[41] M. A. A. F. d. C. T. Carrondo, M. Coll, J. Aymami, A. H. J. Wang G. A. V. Marel, J. H. V. Boom, A. Rich, Binding of a Hoechst Dye to d(CGCGATATCGCG) and Its Influence on the Conformation of the DNA Fragment, Biochemistry. 28 (1989) 7849–7859, doi: 10.1021/bi00445a047.

[42] S. A. Miller, D. R. Gallie, D. E. Sleat, J. W. Watts, P. C. Turner, T. M. Wilson, Mutational analysis of the tobacco mosaic virus 5’-leader for altered ability to enhance translation, Nucleic Acids Res. 16 (1988) 883–893, doi: 10.1093/nar/16.3.883.

[43] N. Spink, D. G. Brown, J. V. Skelly, S. Neidle, Sequence-dependent effects in drug-DNA interaction: the crystal structure of Hoechst 33258 bound to the d(CGCAAATTTGCG) 2 duplex, Nucleic Acids Res. 22 (1994) 1607–1612, http://nar.oxfordjournals.org/.

[44] D. G. Brown, M. R. Sanderson, E. Garman, S. Neidle, Crystal structure of a berenil-d(CGCAAATTTGCG) complex. An example of drug-DNA recognition based on sequence-dependent structural features, J. Mol. Biol. 226 (1992) 481–490, doi: 10.1016/0022-2836(92)90962-J.

[45] D. G. Brown, M. R. Sanderson, J. V. Skelly, T. C. Jenkins, T. Brown, E. Garman, D. I. Stuart, S. Neidle, Crystal structure of a berenil -dodecanucleotide complex: The role of water in sequence-specific ligand binding, EMBO J. 9 (1990) 1329–1334, doi: 10.1002/j.1460-2075.1990.tb08242.x.

[46] S. Borghini, M. Vargiolu, M. D. Duca, R. Ravazzolo, I. Ceccherini, Nuclear factor Y drives basal transcription of the human TLX3, a gene overexpressed in T-cell acute lymphocytic leukemia, Mol. Cancer Res. 4 (2006) 635–643, doi: 10.1158/1541-7786.MCR-05-0250.

[47] T. Boulikas, Common structural features of replication origins in all life forms, J. Cell. Biochem. 60 (1996) 297–316, doi: 10.1002/(SICI)1097-4644(19960301)60:3<297::AID-JCB2>3.0.CO;2-R.

[48] R. E. Holmlin, E. D. A. Stemp, J. K. Barton, Ru(phen)2dppz^2+^ Luminescence: Dependence on DNA Sequences and Groove-Binding Agents, Inorg Chem. 37 (1998) 9–34, doi: 10.1021/ic970869r.

[49] A. M. Pyle, J. P. Rehmann, R. Meshoyrer, N. J. Turro, J. K. Barton, C. V. Kumar, Mixed-Ligand complexes of ruthenium(II): Factors governing binding to DNA, J. Am. Chem. Soc. 111 (1989) 3051–3058, doi: 10.1021/ja00190a046.

[50] A. S. Abdelhameed, A. H. Bakheit, M. S. Mohamed, W. M. Eldehna, H. A. Abdel-Aziz, M. I. Attia, Synthesis and biophysical insights into the binding of a potent antiproliferative non-symmetric bis-isatin derivative with bovine serum albumin: Spectroscopic and molecular docking approaches, Appl. Sci. 7 (2017) 617, doi: 10.3390/app7060617.

[51] T. D. Goddard, D. G. Kneller, SPARKY 3, University of California, San Francisco.

[52] D. A. Case, R. C. Walker, T. E. Cheatham, et al., Amber 18, Univ. California, San Fr. (2018), http://ambermd.org/doc12/Amber18.pdf.

[53] R. Lavery, M. Moakher, J. H. Maddocks, D. Petkeviciute, K. Zakrzewska, Conformational analysis of nucleic acids revisited: Curves+, Nucleic Acids Res. 37 (2009) 5917–5929, doi: 10.1093/nar/gkp608.

[54] M. S. Nair, S. D’Mello, R. Pant, K. M. Poluri, Binding of resveratrol to the minor groove of DNA sequences with AATT and TTAA segments induces differential stability, J. Photochem. Photobiol. B Biol. 170 (2017) 217–224, doi: 10.1016/j.jphotobiol.2017.04.016.

[55] J. M. Lopez-Nicolas, F. Garcia-Carmona, Aggregation state and pKa values of (E)-resveratrol as determined by fluorescence spectroscopy and UV-visible absorption, J. Agric. Food Chem. 56 (2008) 7600–7605, doi: 10.1021/jf800843e.

[56] U. Pal, S. K. Pramanik, B. Bhattacharya, B. Banerji, N. C. Maiti, Binding interaction of a novel fluorophore with serum albumins: steady state fluorescence perturbation and molecular modeling analysis, Springerplus. 4 (2015) 548, doi: 10.1186/s40064-015-1333-8.

[57] G. Lober, The fluorescence of dye-nucleic acid complexes, Journal of Luminescence. 22 (1981) 221–265, doi: 10.1016/0022-2313(81)90022-3.

[58] H. Ihmels, K. Faulhaber, K. Wissel, G. Viola, D. Vedaldi, 6-aminoacridizinium bromide: A fluorescence probe which lights up in AT-rich regions of DNA, Org. Biomol. Chem. 1 (2003) 2999–3001, doi: 10.1039/b305439j.

[59] J. Kypr, I. Kejnovska, D. Renciuk, M. Vorlickova, Circular dichroism and conformational polymorphism of DNA, Nucleic Acids Res. 37 (2009) 1713–1725, doi: 10.1093/nar/gkp026.

[60] J. B. Chaires, Energetics of drug–DNA interactions, Biopolymers. 44 (1997) 201–215, doi: 10.1002/(SICI)1097-0282(1997)44:3<201::AID-BIP2>3.0.CO;2-Z.

[61] J. B. Chaires, A thermodynamic signature for drug-DNA binding mode, Arch. Biochem. Biophys. 453 (2006) 26–31, doi: 10.1016/j.abb.2006.03.027.

[62] J J. G. Duguid, V. A. Bloomfield, J. M. Benevides, G. J. Thomas, DNA Melting Investigated by Differential Scanning Calorimetry and Raman Spectroscopy, Biophysical Journal. 71 (1996) 3350–3360, doi: 10.1016/S0006-3495(96)79528-0.

[63] K. J. Breslauer, E. Freire, M. Straume, Calorimetry: A tool for DNA and ligand-DNA studies, Methods Enzymol. 211 (1992) 533–567, doi: 10.1016/0076-6879(92)11030-M.

[64] G. M. Clore, A. M. Gronenborn, D. S. Moss, I. J. Tickle, Refinement of the solution structure of the B DNA hexamer 5’d(C-G-T-A-C-G)2 on the basis of inter-proton distance data, J. Mol. Biol. 185 (1985) 219–226, doi: 10.1016/0022-2836(85)90195-0.

[65] R. V. Hosur, G. Govil, H. T. Miles, Application of two-dimensional NMR spectroscopy in the determination of solution conformation of nucleic acids, Magn. Reson. Chem. 26 (1988) 927–944, doi: 10.1002/mrc.1260261019.

[66] K. Wüthrich, NMR with Proteins and Nucleic Acids, Europhys. News. 17 (1986) 11–13, doi: 10.1051/epn/19861701011.

[67] S. E. Pineda-Sanabria, I. M. Robertson, B. D. Sykes, Structure of trans-resveratrol in complex with the cardiac regulatory protein troponin C, Biochemistry. 50 (2011) 1309–1320, doi: 10.1021/bi101985j.

[68] D. Koh, K. H. Park, J. Jung, H. Yang, K. H. Mok, Y. Lim, Complete assignment of the ^1^H and ^13^C NMR spectra of resveratrol derivatives, Magn. Reson. Chem. 39 (2001) 768–770, doi: 10.1002/mrc.910.

[69] H. S. Park, E. H. Kim, M. R. Kang, I. K. Chung, C. Cheong, W. Lee, Spectroscopic studies on interaction of protoberberines with the deoxyoligonucleotide d(GCCGTCGTTTTACA)2, Bull. Korean Chem. Soc. 25 (2004) 1559–1563, doi: 10.5012/bkcs.2004.25.10.1559.

[70] S. Mazzini, M. C. Bellucci, R. Mondelli, Mode of binding of the cytotoxic alkaloid berberine with the double helix oligonucleotide d(AAGAATTCTT)2, Bioorganic Med. Chem. 11 (2002) 505–514, doi: 10.1016/S0968-0896(02)00466-2.

[71] D. G. Gorenstein, D. Kar, Effect of Bond Angle Distortion on Torsional Potentials. Ab Initio and CNDO/2 Calculations on Dimethoxymethane and Dimethyl Phosphate, J. Am. Chem. Soc. 99 (1977) 672–677, doi: 10.1021/ja00445a002.

[72] D. G. Gorenstein, 31P NMR of DNA, Methods Enzymol. 211 (1983) 254–286, 10.1016/0076-6879(92)11016-C.

[73] S. Mazzini, R. Mondelli, E. Ragg, Structure and dynamics of intercalation complexes of anthracyclines with d(CGATCG)2 and d(CGTACG)2. 2D-^1^H and ^31^P NMR investigations, J. Chem. Soc. Perkin Trans. 2 (1998) 1983–1991, doi.org/10.1039/A802350F.

[74] M. S. Searle, J. G. Hall, W. A. Denny, L. P. G. Wakelin, MR Studies of the Interaction of the Antibiotic Nogalamycin with the Hexadeoxyribonucleotide Duplex d(5’-GCATGC)2, Biochemistry. 27 (1988) 4340–4349, doi: 10.1021/bi00412a022.

[75] A. Favier, M. Blackledge, J. P. Simorre, S. Crouzy, V. Dabouis, A. Gueiffier, D. Marion, J. C. Debouzy, olution Structure of 2-(Pyrido[1,2-e]purin-4-yl)amino-ethanol Intercalated in the DNA Duplex d(CGATCG)2, Biochemistry. 40 (2001) 8717–8726, doi: 10.1021/bi0024963.

[76] G G. Kotovych, J. William Lown, J. P. K. Tong, High-field^1^H and ^31^P NMR studies on the binding of the anticancer agent mitoxantrone to d[CpGpApTpCpG]2, J. Biomol. Struct. Dyn. 4 (1986) 111–124, doi: 10.1080/07391102.1986.10507649.

[77] S. Mazzini, M. C. Bellucci, S. Dallavalle, F. Fraternali, R. Mondelli, Mode of binding of camptothecins to double helix oligonucleotides, Org. Biomol. Chem. 2 (2004) 505–513, doi: 10.1039/b312780j.

[78] T. W. Plumbridge, J. R. Brown, Spectrophotometric and fluorescence polarization studies of the binding of ethidium, daunomycin and mepacrine to DNA and to poly(I · C), Biochimica et Biophysica Acta. 479 (1977) 441–449, doi: 10.1016/0005-2787(77)90037-5.

[79] J. B. Chaires, N. Dattagupta, D. M. Crothers, Self-Association of Daunomycin, Biochemistry. 21 (1982) 3933–3940, doi: 10.1021/bi00260a004.

[80] D. J. Patel, Structure and Dynamics of Poly(dG-dC) in Solution Steroid Diamine•Nucleic Acid Complexes and Generation of an ‘Alternating B-DNA’ Conformation in High Salt, Stereodynamics of Molecular Systems. (1979) 397–422, doi.org/10.1016/B978-0-08-024629-1.50031-9.

[81] I. Ivani, P. Dans, A. Noy, et al., Parmbsc1: a refined force field for DNA simulations, Nat. Methods. 13 (2016) 55–58, doi: 10.1038/nmeth.3658.

[82] J. Wang, R. M. Wolf, J. W. Caldwell, P. A. Kollman, D. A. Case, Development and testing of a general amber force field, J. Comput. Chem. 25 (2004) 1157–1174, doi: 10.1002/jcc.20035.

[83] W. Jorgensen, J. Chandrasekhar, J. Madura, R. Impey, M. Klein, Comparison of Simple Potential Functions for Simulating Liquid Water, J. Chem. Phys. 79 (1983) 926–935, doi: 10.1063/1.445869.

[84] D. R. Roe T. E. Cheatham, PTRAJ and CPPTRAJ: Software for Processing and Analysis of Molecular Dynamics Trajectory Data, J. Chem. Theory Comput. 9 (2013) 3084–3095, doi: 10.1021/ct400341p.

[85] A. A. Wood, C. M. Nunn, A. Czarny, D. W. Boykin, S. Neidle, Variability in DNA minor groove width recognized by ligand binding: The crystal structure of a bis-benzimidazole compound bound to the DNA duplex d(CGCGAATTCGCG)2, Nucleic Acids Res. 23 (1995) 3678–3684, doi: 10.1093/nar/23.18.3678.

[86] C. A. Bewley, A. M. Gronenborn, G. M. Clore, Minor groove-binding architectural proteins: structure, function, and DNA recognition, Annu. Rev. Biophys. Biomol. Struct. 27 (1998) 105–131, doi: 10.1146/annurev.biophys.27.1.105.

[87] C. H. Ríos Martínez, L. Lagartera, C. Trujillo, C. Dardonville, Bisimidazoline arylamides binding to the DNA minor groove: N1-hydroxylation enhances binding affinity and selectivity to AATT sites, Medchemcomm. 6 (2015) 2036–2042, doi: 10.1039/C5MD00292C.

[88] M. Rettig, M. W. Germann, S. Wang, W. D. Wilson, Molecular Basis for Sequence-Dependent Induced DNA Bending, ChemBioChem 14 (2013) 323–331, doi: 10.1002/cbic.201200706.

