## Supplementary material for "Exploring the binding of resveratrol to an oncogene promoter DNA sequence d(CCAATTGG)_2_ through multispectroscopic, nuclear magnetic resonance and molecular dynamics studies"

**Supplementary material Table 1A. Chemical shift values (ppm) of d-(CCAATTGG)$_2$ in 90%H$_2$0 +10%D$_2$0 at 298K.**

| PROTONS | C1 | | | C2 | | | A3 | | | A4 | | |
|---|---|---|---|---|---|---|---|---|---|---|---|---|
| | δb | δf | **Δ δ** | δb | δf | **Δ δ** | δb | δf | **Δ δ** | δb | Δf | **Δ δ** |
| H8/H6 | 7.75 | 7.77 | -0.02 | 7.61 | 7.61 | 0 | 8.32 | 8.31 | 0.01 | 8.28 | 8.22 | 0.06 |
| H1’ | 5.99 | 5.99 | 0 | 5.27 | 5.27 | 0 | 6.01 | 6.01 | 0 | 6.22 | 6.21 | 0.01 |
| H2’ | 2.05 | 2.05 | 0 | 2.12 | 2.12 | 0 | 2.86 | 2.86 | 0 | 2.65 | 2.65 | 0 |
| H2’’ | 2.49 | 2.49 | 0 | 2.38 | 2.39 | -0.01 | 2.96 | 2.97 | -0.01 | 2.94 | 2.94 | 0 |
| H3’ | 4.62 | 4.67 | -0.05 | | | | | | | 5.03 | 5.09 | -0.06 |
| H4’ | 4.1 | 4.11 | -0.01 | 4.06 | 4.06 | 0 | 4.45 | 4.46 | -0.01 | 4.5 | 4.52 | -0.02 |
| H5’ | 3.9 | 3.89 | 0.01 | | | | 4.26 | 4.29 | -0.03 | 4.268 | 4.301 | -0.033 |
| H5’’ | 3.761 | 3.76 | 0.001 | | | | | | | 4.147 | 4.174 | -0.027 |
| H5/H2/CH3 | 5.97 | 5.99 | -0.02 | 5.73 | 5.73 | 0 | 7.71 | 7.68 | 0.03 | 7.71 | 7.68 | 0.03 |
| NH2b | 8.08 | 8.04 | 0.04 | 8.68 | 8.7 | -0.02 | 7.32 | 7.33 | -0.01 | 7.24 | 7.26 | -0.02 |
| NH2nb | 7.24 | 7.22 | 0.02 | 6.92 | 6.95 | -0.03 | 6.36 | 6.4 | -0.04 | 6.21 | 6.2 | 0.01 |
| NH | NA | NA | NA | NA | NA | NA | NA | NA | NA | NA | NA | NA |
| **Protons** | **T5** | | | **T6** | | | **G7** | | | **G8** | | |
| | δb | δf | Δ δ | δb | δf | Δ δ | δb | δf | Δ δ | δb | Δf | Δ δ |
| H8/H6 | 7.12 | 7.15 | -0.03 | 7.24 | 7.22 | 0.02 | 7.65 | 7.83 | -0.18 | 7.8 | 7.79 | 0.01 |
| H1’ | 5.94 | 5.96 | -0.02 | 5.84 | 5.88 | -0.04 | 5.64 | 5.67 | -0.03 | 6.11 | 6.13 | -0.02 |
| H2’ | 1.96 | 1.96 | 0 | 1.96 | 1.96 | 0 | 2.58 | 2.54 | 0.04 | 2.33 | 2.33 | 0 |
| H2’’ | 2.49 | 2.49 | 0 | 2.31 | 2.33 | -0.02 | 2.68 | 2.68 | 0 | 2.55 | 2.55 | 0 |
| H3' | | 4.85 | | | | | | | | 4.66 | 4.67 | -0.01 |
| H4' | | 4.35 | | 4.1 | 4.11 | -0.01 | 4.35 | 4.35 | 0 | 4.22 | 4.22 | 0 |
| H5' | | | | 4.14 | 4.15 | -0.01 | | | | 4.21 | | |
| H5" | | | | 4.04 | 4.10 | -0.06 | | | | 4.17 | | |
| H5/H2/CH3 | 1.33 | 1.34 | -0.01 | 1.59 | 1.61 | -0.02 | NA | NA | NA | NA | NA | NA |
| NH2b | NA | NA | NA | NA | NA | NA | 7.98 | 7.97 | 0.01 | 7.50 | 7.49 | -0.01 |
| NH2nb | NA | NA | NA | NA | NA | NA | 6.38 | 6.4 | -0.02 | 6.78 | 6.77 | 0.01 |
| NH | 13.75 | 13.76 | -0.01 | 13.99 | 14 | -0.01 | 12.91 | 12.89 | 0.02 | NA | NA | NA |

δb = Chemical shift values of bound form, δf =Chemical shift values of free form, Δδ = $\delta_b$ - $\delta_{f,}$ -ve Δδ indicates upfield shift, +ve Δδ indicates downfield shift.

**Supplementary material Table 1B. Intermolecular cross peaks between resveratrol and d-(CCAATTGG)$_2$.**

| Resveratrol- DNA cross peaks | Calculated NOE distances (Å) |
|---|---|
| RES 2,6 H- A3NH$_2^{nb}$ | 3.33 |
| RES 2,6 H -T6H6 | 3.32 |
| RES 2,6 H –A3/A4 H2 | -- |

**Supplementary material Table 1C. Intramolecular cross peaks within resveratrol, observed in complex spectrum.**

| **Intramolecular Resveratrol cross peaks** | |
|---|---|
| H2′/H6′ | Hα′ |
| H2′/H6′ | H3′/ H5′ |
| H2′/H6′ | Hα |

**Supplementary material Table 2. Chemical Shifts of resveratrol protons in deuterated ethanol and $D_20$.**

| Resveratrol Protons | Chemical shift(ppm) in ethanol | Chemical Shift(ppm) in $D_20$. |
|---|---|---|
| 2' ,6' | 7.31 | 7.51 |
| 3' ,5' | 6.79 | 6.91 |
| α' | 6.79 | 7.17 |
| α | 6.76 | 6.98 |
| 2 ,6 | 6.47 | 6.67 |
| 4 | 6.21 | 6.32 |

**Supplementary material Table 3.** Chemical shift of $^{31}P$ resonances (ppm) of DNA octamer in d-(CCAATTGG)$_2$complexed with resveratrol at 298K

| | $\delta_f$ | $\delta_b$ | $\Delta\delta = \delta_b - \delta_f$ |
|---|---|---|---|
| C1pC2 | -0.785 | -0.782 | 0.003 |
| C2pA3 | -0.425 | -0.426 | -0.001 |
| A3pA4 | -1.016 | -1.033 | -0.017 |
| A4pT5 | -1.144 | -1.157 | -0.013 |
| T5pT6 | -1.144 | -1.157 | -0.013 |
| T6pG7 | -0.855 | -0.908 | -0.053 |
| G7pG8 | -0.522 | -0.519 | 0.003 |

δb = Chemical shift values of bound form, δf =Chemical shift values of free form, $\Delta\delta = \delta_b - \delta_f$, -ve, Δδ indicates upfield shift, +ve Δδ indicates downfield shift

**Supplementary material Fig.1A**. NOE connectivities of resveratrol-DNA complex showing base-H2′-H2′′ region.

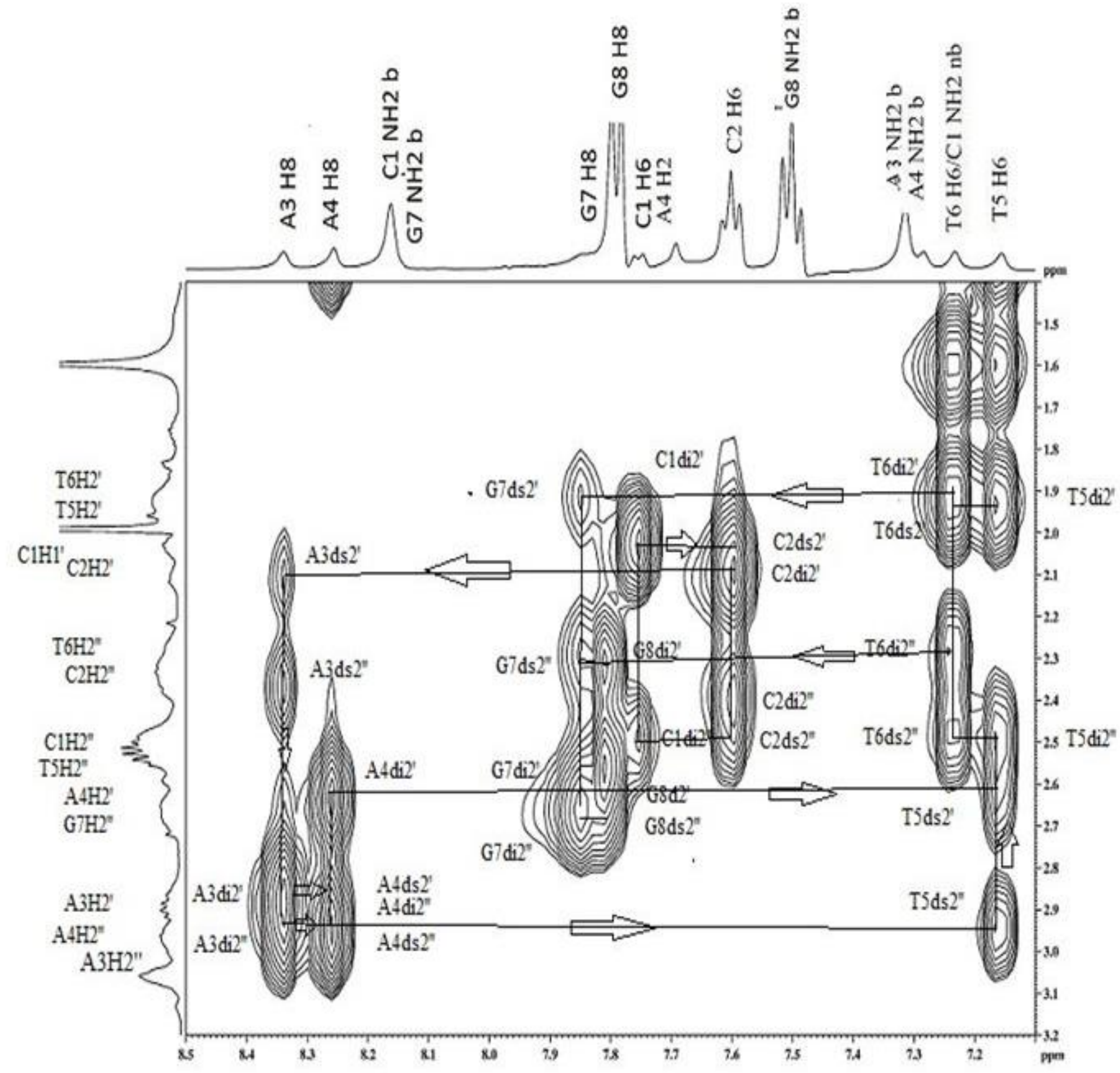

**Supplementary material Fig.1B**. NOE connectivities of resveratrol-DNA complex showing base-NH region.

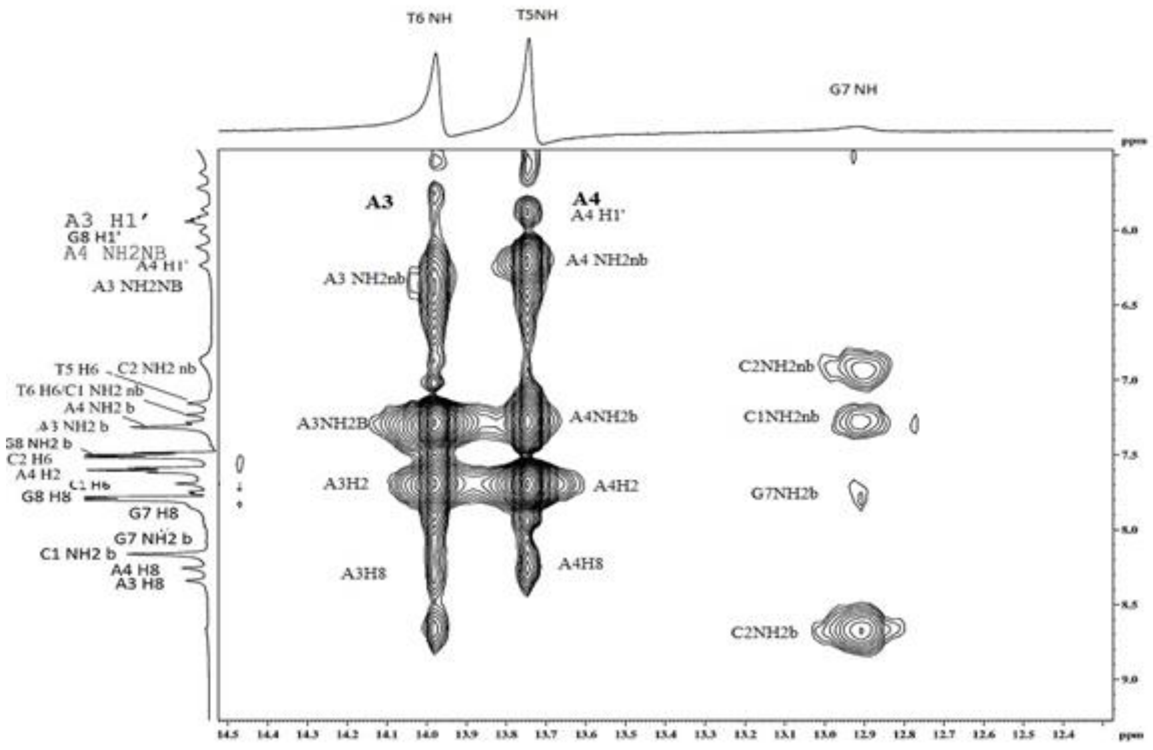
